# Non-equilibrium conditions inside rock pores drive fission, maintenance and selection of coacervate protocells

**DOI:** 10.1101/2021.07.08.451414

**Authors:** Alan Ianeselli, Damla Tetiker, Julian Stein, Alexandra Kühnlein, Christof Mast, Dieter Braun, T-Y Dora Tang

## Abstract

Key requirements for the first cells on Earth include the ability to compartmentalize and evolve. Compartmentalization spatially localizes biomolecules from a dilute pool and an evolving cell which grows and divides permits mixing and propagation of information to daughter cells. Complex coacervate microdroplets are excellent candidates as primordial cells with the ability to partition and concentrate molecules into their core and support primitive and complex biochemical reactions. However, the evolution of coacervate protocells by fusion, growth and fission has not yet been demonstrated. In this work, a primordial environment initiated the evolution of coacervate-based protocells. Gas bubbles inside heated rock pores perturb the coacervate protocell distribution and drive the growth, fusion, division and selection of coacervate microdroplets. This setting provides a primordial non-equilibrium environment. Our findings describe how common gas bubbles within heated rock pores induce the early evolution processes of coacervate-based protocells, providing a compelling scenario for the evolution of membrane-free coacervate microdroplets on the early Earth.

## Introduction

Compartmentalization is a key feature of modern biological systems and has been hypothesized to play an important role during the origin of life by spatially localizing molecules and facilitating the first chemical reactions^1,2^. One viable route to compartmentalization is via liquid-liquid phase separation of oppositely charged polyelectrolytes in aqueous solution^3^. This process leads to the formation of membrane-free chemically enriched droplets. These coacervate microdroplets are intriguing protocell models as they form with little chemical identity under a broad range of physico-chemical conditions^4^; they localize and concentrate a range of different molecules^5–7^ and exhibit molecular selectivity by partitioning^8–10^. In addition, coacervate droplets facilitate the assembly of fatty acid bilayers on their outer surface^11^ and readily support catalytic reactions such as primitive RNA catalysis^12,13,14^. This provides a pathway to membrane bound compartmentalization as observed in modern biology and a connection to the RNA-peptide world hypothesis.

Fusion events, division and maintenance of coacervate protocells would have been essential for the evolution of compartmentalized molecules. Fusion and growth of protocells are necessary for the exchange of molecules and genetic material^15^ and it has been shown that the incorporation of free components by direct fusion with other protocells^16^ or by external electric fields^17^ can be achieved in a laboratory setting. In solution, these coacervate droplets will tend to coalescence eventually forming a coacervate bulk macrophase^18,19^ which limits their role as protocells. The division of coacervate protocells is required to transfer molecular information to succeeding daughter protocells that can pass evolutionary advantages to the next generation. To achieve division, modern cells make use of a complex machinery of regulatory proteins, scaffold proteins, enzymes and chemical messengers^20^. In the prebiotic world, division must have relied on other factors. Some studies suggest that division of lipid-based vesicles can be triggered by osmotic changes^21^, chemical changes^22^, temperature^23^ and shearing forces^24^. In comparison, less is known about the division mechanisms of membrane-free coacervate-based protocells which are chemically enriched. One theoretical study predicts that budding of chemically active membrane free droplets is achieved by the flux of substrate and product across the interface which lies in a particular surface tension regime^25^. Despite this prediction, there has been no experimental realization of fission of membrane-free protocells with or without chemical input. Furthermore, it has still not experimentally shown how they would behave under prebiotically plausible non-equilibrium conditions.

To this end, pores in a thermal gradient provide a unique, facile and prebiotically feasible route to perturbing the system away from it’s equilibria^26^. Here, capillary flows induced by heat fluxes within millimeter-sized pores have been shown to accumulate molecules based on their size at the gas-water interface of gas inclusions. Simulations and experiments show that there are two main forces acting at the interface: capillary flows from the cold to the warm side and perturbative fluxes after the precipitation of water^27,28^. These forces induced rapid movements of particles, driving their contact and fusion. Under these conditions, lipid molecules accumulate at the interface to create vesicular structures and undergo fission driven by Marangoni flows and convection. These previous studies indicate that the growth, division and maintenance of coacervate droplets could be manipulated by the physical flows within thermal pores.

Herein, we study the effect of out-of-equilibrium conditions provided by heated pores containing gas bubbles, a common primordial scenario^26^, on the growth and division mechanisms of complex coacervate microdroplets formed by mixing polyanionic (carboxymethyl dextran (CM-Dex), adenosine 5’-triphosphate (ATP)) and polycationic (poly-diallyl dimethylammonium (PDDA), poly-L-lysine (pLys)) species. Even though the coacervates in this study might not be generated from prebiotically relevant molecules they provide a robust model system for reconciling the general role of heat-induced out-of-equilibrium systems on coacervate microdroplets.

We show that the accumulation of coacervate forming components at the gas-water interface of the gas bubble^28^ drives growth by fusion of the coacervate microdroplets. Droplets of up to 300 μm in size are formed and maintained over time. This property is not observed under equilibrium conditions where droplets coalesce to eventually form a single coacervate macrophase (Figure S5.1)^18,19^. intriguingly, the microfluidic water cycle induced by the thermal gradient^27^ creates perturbative fluxes at the gas-water interface that lead to the fission and fragmentation of the coacervate droplets using purely physical processes (Figure 1a-c). This offers direct evidence that physical forces within a confined environment are sufficient to provide the mechanism of membrane-free protocell division without complex machinery or targeted chemical reactions. Furthermore, the environment provided the ability to create and select for separate populations of droplets with different chemical composition. Specifically, the out-of-equilibrium conditions were able to overcome the intrinsic preference of RNA to coacervate with pLys^29^, yielding RNA:pLys droplets also enriched with CM-Dex at the gas-water interface. in the bulk, the coacervate droplets were formed mainly by RNA and pLys. This means, the thermal gradient in combination with the gas bubble led to the creation and spatial segregation of two different populations of coacervate droplets with different composition: oligonucleotide:poly-peptide (RNA:pLys) coacervate droplets in the bulk and sugar:oligonucleotide:poly-peptide (CM-Dex:RNA:pLys) droplets at the gas-water interface.

**Figure 1.**
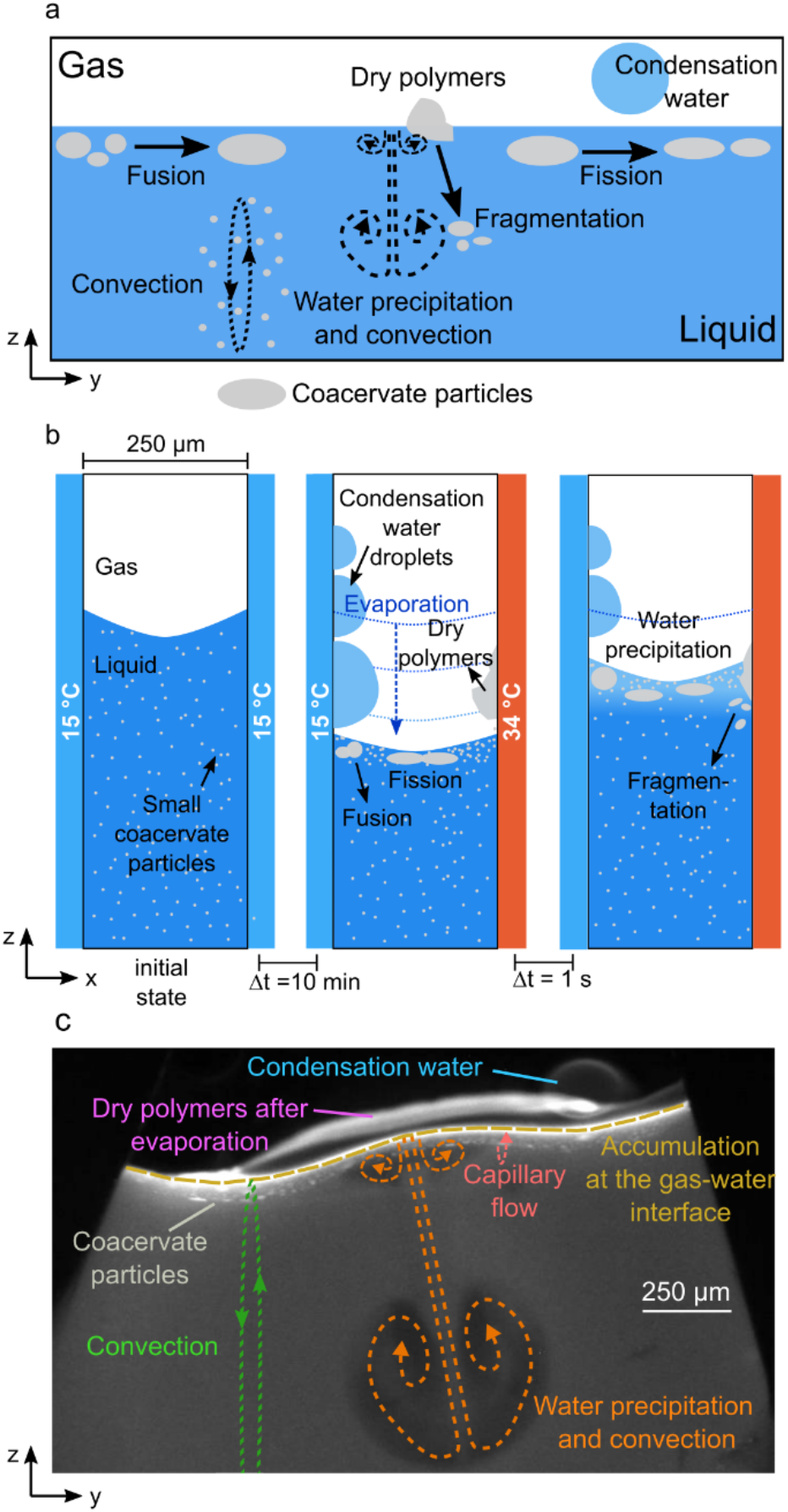
Fusion, division and transport of coacervate protocells inside a thermal pore. a) Scheme of coacervate transport, accumulation, growth and division at the gas-water interface. b) (Left) no heating: pre-formed small coacervate droplets in the bulk. (Center) temperature gradient: the droplets accumulate and fusion and fission are observed. (Right) water precipitation drives coacervate fragmentation. c) Fluorescence image showing evaporation, water condensation, wet-dry cycles, convection and capillary flows at the gas-water interface of the thermal pore. Conditions for c) were: CM-Dex:PDDA total polymer concentration 2mM (molar ratio 6:1, [carboxy]/[amine] = 5) + 0.1% FITC-labeled CM-Dex, 10 mM MgCl_2_, 10 mM Tris pH 8, temperature gradient of 19 °C (hot side 34 °C, cold side 15 °C).

We present the proposed mechanisms as a prebiotic model for membrane-free protocell growth, division, and evolution, since the only requirements are simple and ubiquitous physical conditions that could be found inside heated rock pores on the early Earth.

## Results

### The gas-water interface accumulates coacervate droplets and facilitates fusion

To characterize the effect of non-equilibrium perturbations on coacervate microdroplets, we experimentally recreated a heated rock pore filled with liquid and gas bubbles as described previously^27,28^. In brief, a PTFE sheet (250 μm thick) cut with sharp triangular structures was placed between an optically transparent sapphire and a silica plate (Figure 2a). Liquids were loaded into the chamber through microfluidic tubings and gas bubbles were created by incomplete filling of the liquid into the triangular cavities (Figure 2b). The sample chamber was loaded onto a custom-built microscope (see Materials and Methods and supplementary section 1) and a temperature gradient was generated by differentially heating the sapphire with rod resistors inserted into a copper holder, and cooling the copper back plate through a connection to a water bath (Figure 2c-d). The temperature gradients were varied between 15 °C to 29 °C with an accuracy of ± 1 °C. Imaging was provided through the transparent sapphire with the camera focused on the cold wall. This chamber is also referred to as a “thermal trap”.

**Figure 2.**
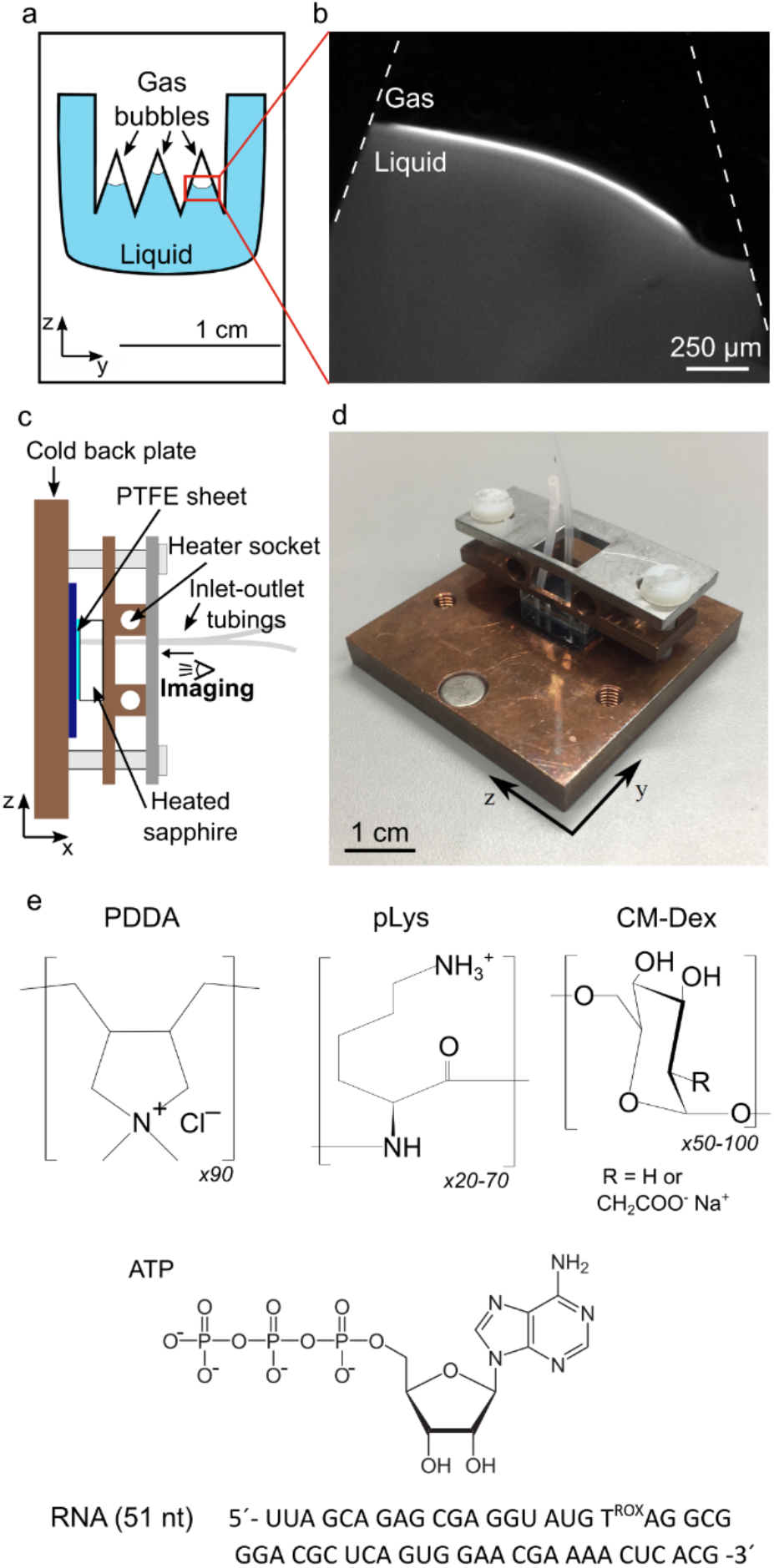
Scheme of the thermal trap used in the experiments. a) Scheme of the PTFE interspacer. The triangular structures cause the formation of gas bubbles. b) Fluorescence image of the gas bubble in a temperature gradient (CM-Dex:PDDA 6:1 molar ratio, [carboxy]/[amine] = 5 + 0.1% FITC-labeled CM-Dex, total conc 2 mM). Lateral sketch (c) and photo (d) of the thermal trap. e) Chemical structure of the components used: PDDA (poly-diallyl dimethylammonium chloride), pLys (poly-L-lysine), CM-Dex (carboxymethyl-dextran), ATP (adenosine triphosphate) RNA sequence (51 nt).

Coacervate microdroplet dispersions were prepared by mixing negatively charged modified sugars carboxymethylated-dextran (CM-Dex, degree of polymerization between 50-100, with 1 carboxyl group every 3 repeats) or adenosine triphosphate (ATP) with positively charged polyelectrolytes, either poly-L-lysine (pLys, degree of polymerization of 20 to 70) or polydiallydimethylammonium chloride (PDDA, degree of polymerization of 90) (Figure 2e). CM-Dex:PDDA and CM-Dex:pLys mixtures were prepared at molar ratios of 6:1 and 4:1, respectively, whilst ATP:PDDA and ATP:pLys droplets were prepared at 4:1 molar ratio. The molar ratios correspond to a [carboxyl] to [amine] ratio of 5 (CM-Dex:PDDA) or 7 (CM-Dex:pLys). Such ratios were optimized in previous works to yield a good amount of coacervation^30,31^. The total polymer concentrations were varied between 2 and 20 mM. The starting concentration dictated the density of coacervate droplets within the dispersion and the final amount of material accumulated at the gas-water interface. In order to visualize the coacervate droplets, we added 0.1% FITC labelled CM-Dex or pLys. The coacervate dispersions were prepared in either 0.1 M Na^+^ bicine buffer pH 8.5, or 10 mM Tris (pH 8) and 4 mM MgCl_2_. Control experiments showed that there was no appreciable difference between the two different buffers regarding the dynamics of the coacervate within the thermal trap (see supplementary section 2). Therefore, we used both buffers interchangeably throughout our experiments to highlight the generality of our findings.

Upon loading the coacervate dispersion (20 mM CM-Dex:PDDA in 0.1 M Na^+^ bicine buffer, pH 8.5) into the thermal trap, microscopy images (taken every ~ 1 second) showed the presence of small coacervate droplets (< 10 μm) evenly dispersed throughout the chamber (see Figure 3a). After differential heating at the two sides of the trap (warm side 49 °C, cold side 20 °C), the fluorescent droplets experienced convective flows in the bulk of the solution. The speed of the convective flow could be modulated by the temperature difference as observed in previous simulations^27^. Interestingly, we saw that the coacervate droplets in the bulk solution were transported by the convection flow to the gas-water interface where they accumulated and started growing by fusion (Figure 3b-c and supplementary movie 1). At the interface the droplets moved parallel to the interface driving contact and coalescence events. An individual fusion process between two coacervate droplets required a few seconds (from 1 to 10 seconds) and resulted in elliptically-shaped coacervates. Figure 3d shows the process of fusion between 3 large coacervate droplets.

**Figure 3.**
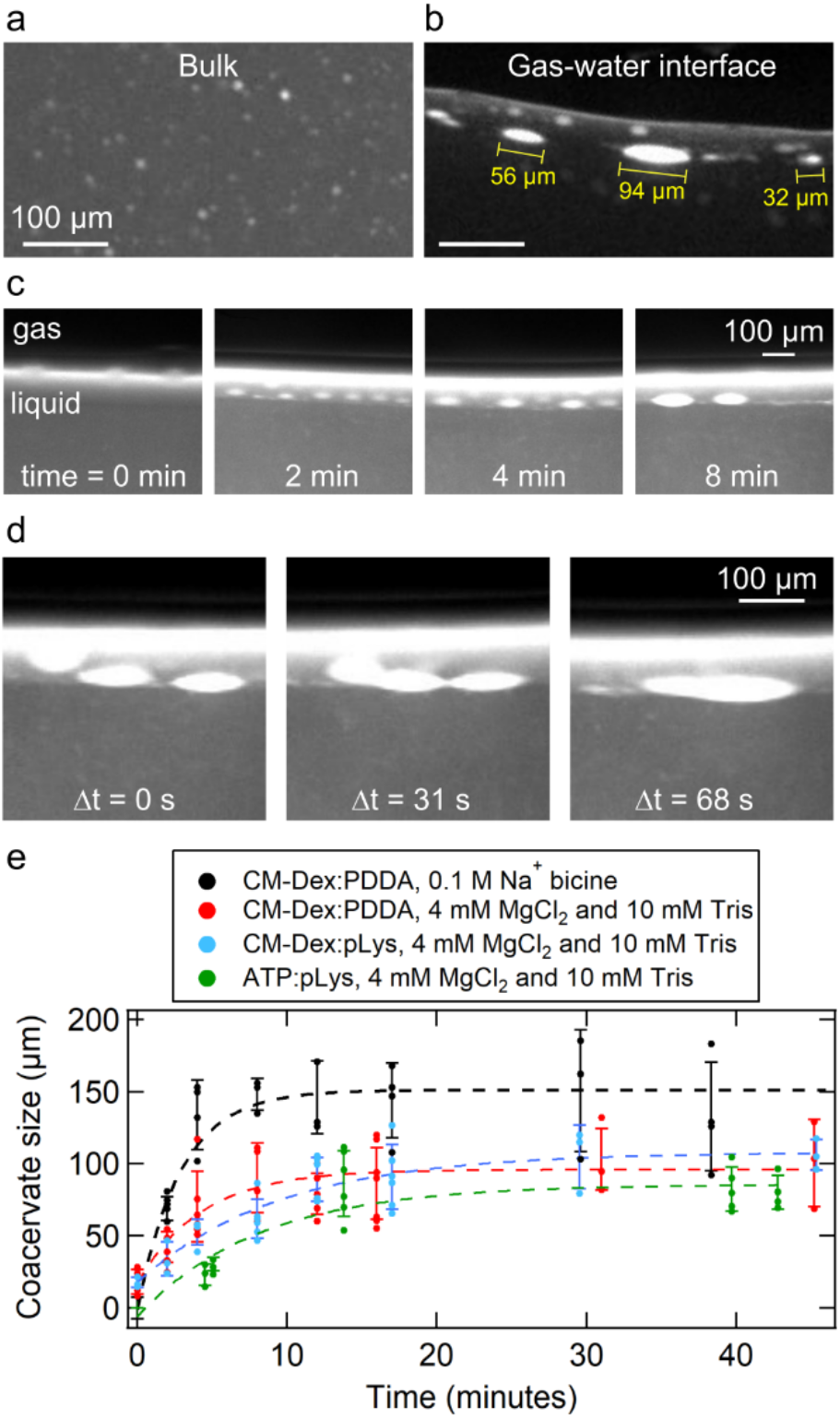
Coacervate droplets accumulate and fuse at the gas-water interface. a) Fluorescent microscopy images of coacervate droplets in the bulk (a) and at the gas-water interface (b). c) Coacervate droplets at the interface after different times in a thermal gradient (t=0 min, 2 min, 4 min and 8 min). d) Microscopy images showing a fusion event between three coacervates. e) Quantification of coacervate size over time for different buffer and coacervate compositions. Each data point represents the mean and standard deviation of approximately 5 different larger droplets at the gas-water interface. The dashed lines represent a phenomenological exponential fit.

The growth of the coacervates over time was quantified from the optical microscopy images. Using LabVIEW, the average horizontal size was measured at different times (as depicted in Figure 3b). Analysis of the CM-Dex:PDDA coacervates reached a maximal average size of 150 μm. Experiments with a different buffer (10 mM Tris pH 8 and 4 mM MgCl_2_) or different polymers of different molecular weights (CM-Dex:pLys, ATP:pLys, ATP:PDDA or CM-Dex:pLys of higher molecular weight) showed comparable behavior with minor differences on the final coacervate size (Figure 3e and supplementary section 2). Note that in our analysis we only measured the horizontal size and not the whole volume of the coacervate droplet. Therefore, we believe that our method was not sensitive to small changes in size. This could be why there was no particular observable effect of the buffer or coacervate type on the final droplet size. However, the method was successful in calculating the average size distribution, as shown in Figure 3e and 5j.

**Figure 4.**
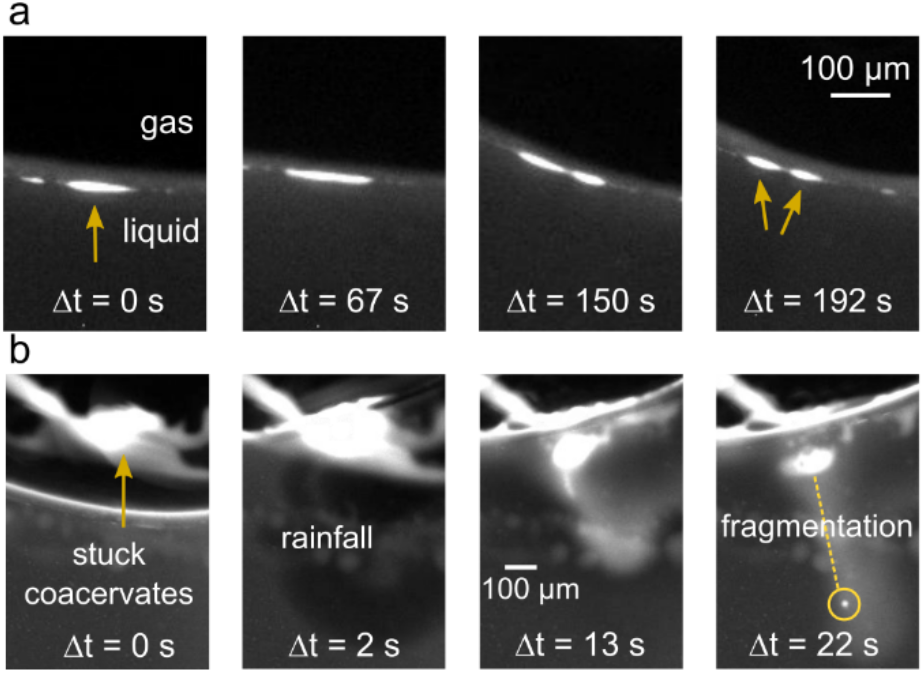
Fission of coacervates induced by (a) interfacial forces and (b) fluxes caused by water precipitation. a) Fission of a coacervate droplet into two smaller droplets, induced by interfacial forces at the gas-liquid interface. The initial droplet (yellow arrow) is slowly stretched (over a time frame of minutes) at the interface until it divides into two smaller droplets. b) Rehydration of stuck coacervates can induce fission by fragmentation, due to the perturbative fluxes caused by precipitating water. It induces a fast mixing of the dry polymers that eventually fragment.

In addition, we characterized the effect of total polymer concentration on the growth rate and the final size of the coacervate droplets by performing a series of experiments with a constant thermal gradient (hot side 49 °C and cold side 20 °C), buffer conditions (4 mM MgCl_2_, 10 mM Tris, pH 8.0) and polymers (CM-Dex:PDDA molar ratio 6:1, [carboxyl]/[amine] = 5), doped with 0.1 % FITC-labeled CM-Dex. The total polymer concentration was varied between 1 and 20 mM (a common concentration range that was used in other studies^12,18,32,33^). Immediately after inserting the coacervate solution in the thermal trap (< 1 min), fluorescent microscopy images were taken every ~ 1 second. The images were analyzed with ImageJ or LabVIEW to determine the growth rate and the average droplet size at steady state (after 1h of thermal gradient) at the interface (supplementary section 3). The final droplet size and the growth rate did not seem to be significantly affected by the initial polymer concentration.

In addition, we noticed that the variability in the size of the droplets between the experiments was large. This could be attributed to oscillatory salt fluctuations induced by the microscale water cycle, together with the intrinsic stochastic nature of droplet fusion. The salt fluctuations induced by microscale water cycles in our thermal trap were previously characterized and showed periodic salt oscillations and perturbative flows caused by water precipitation^27^. While salts are known to have a major impact on coacervation^34,35^, the effects of the salt oscillations on the coacervate droplets in the thermal trap do not appear to adversely affect the droplet stability, as the droplets stay intact at the interface. It is possible that the small fluctuations in salt concentration at the interface can induce local changes in the droplets’ surface charge, influencing droplet fusion and droplet composition. However, it is clear that the droplets are stable under these salt conditions. We estimated a ~1% change in the bulk salt concentration accounting for total volume within the pore vs the volume of water that takes part in precipitation. Therefore, the high variability in droplets’ size and composition that we observed during our analysis was likely due to variations in salt concentrations and the intrinsic stochastic nature of droplet fusion.

Despite this, in all instances (more than 50 different experiments that explored different coacervate conditions, starting concentrations and buffer conditions) we saw that the coacervate droplets accumulated and fused together, indicating that the accumulation, fusion and maintenance of the coacervate droplet at the gas water interface are general phenomena driven by the forces in the thermal trap rather than the chemistry of the coacervate dispersion.

We also performed experiments with starting polymer concentrations below the critical coacervate concentration (CCC), circa 1 mM for the CM-Dex:PDDA coacervate dispersions. At a starting concentration of 0.2 or 0.05 mM, no coacervate droplets were observed using optical microscopy within the resolution of our experiment, despite evident polymer up-concentration at the gas-water interface (supplementary section 3). Our results indicate that the thermal pore acts at the mechanical level to drive fusion of previously existing coacervate droplets followed by droplet division by stretching or fragmentation, and aggregation by wet-dry cycling.

We then wanted to verify that these observed phenomena were attributed to the gas-water interface in combination with thermal flows. To this end, we undertook two control experiments. The first determined the effect of convective flow alone i.e. In the absence of a gas bubble on the coacervate droplets. To do this, coacervate dispersions (CM-Dex:pLys 2 mM ratio 4:1, [carboxyl]/[amine] = 7, 10 mM Tris pH 8.0, 4 mM MgCl_2_) were loaded into a thermal trap without gas bubbles (hot side 49 °C, cold side 20 °C). Time-resolved optical microscopy images showed that the bulk coacervate droplets (<15 μm) were transported in the bulk by the convection flow at a speed of about 1.6 ± 0.4 μm/s but did not undergo fusion events in the bulk solution or accumulate within the trap (see supplementary section 4). We then characterized the behavior of coacervate droplets within the thermal chamber in the absence of thermal flow. At isothermal conditions, almost 100% of coacervate droplets within the pore slowly sedimented to the bottom of the microfluidic chamber where the droplets fused to form a single coacervate droplet, as expected under isothermal conditions^18,19^. In the presence of the thermal gradient, the convection flow in the bulk prevented the coacervate droplets from sedimenting by maintaining them within the thermophoretic flow. The fraction of droplets that survived sedimentation was proportionally dependent on the thermal gradient. Steeper thermal gradients induced faster convection and prevented the sedimentation of a larger fraction of droplets. Finite element simulations of the sedimentation of the coacervate droplets in a thermal trap with comparable thermal gradients to the experiments showed that droplet sedimentation reached steady state after 5 hrs and was maintained up to 30 hrs (supplementary section 5). In comparison, coacervate droplets at the gas-water interface resided at the interface even with very shallow temperature gradients.

Taken together, our results confirm that the flows at a gas-water interface led to the accumulation of coacervate droplets at the interface, fusion events between the droplets and to the maintenance of the droplets against sedimentation. In the absence of the thermal flow the droplets will sediment to the bottom of the pore. Therefore, the combination of convection and capillary flow at the interface maintained the droplets at the gas-water interface or circulating within the bulk for extended periods of time.

### Droplet division at the gas-water interface

Our data show that the opposing forces at the interface lead to the elongation of the droplets (Figure 3c-d). As an elliptical shape has been associated with the initial stages of vesicle division^36^ we wondered whether the forces in our non-equilibrium setting would be strong enough to drive the elliptical deformation of the membrane-free coacervate droplet into a fission event.

We applied a temperature gradient of 19 °C (15°C-34°C) on a coacervate dispersion of CM-Dex:PDDA (molar ratio 6:1, [carboxyl]/[amine] = 5, total polymer concentration 2 mM, with 10 mM Tris pH 8, 4 mM MgCl_2_) doped with 0.1 % FITC-labeled CM-Dex. Time-resolved optical microscopy images showed that the coacervate droplets accumulated, fused and became elliptically elongated at the gas-water interface (Figure 3c-d). Excitingly, upon accumulation, droplets were progressively stretched along the interface until the droplet divided to produce two daughter protocells of a similar size (Figure 4a, supplementary movie 2). Our results confirm that elliptical deformation of the coacervate droplets at the interface do indeed drive droplet division. Droplet stretching and fission occurred as a consequence of the forces induced by the thermal gradient at the gas-water interface. In additional experiments, CM-Dex:pLys droplets also underwent fission events at the interface indicating that this is a general phenomenon that is driven by the physical forces rather than the chemistry or type of coacervate (supplementary section 6).

In addition to convection and capillary forces at the interface, the presence of a gas bubble creates an environmental water cycle - this hypothetical prebiotic scenario may also have an effect on coacervate behavior and properties. For example, wet-dry cycles can lead to the accumulation, drying and rehydration of molecules at a surface. Previous studies^27,28^ have shown that a heated gas bubble in contact with a cold surface within a thermal trap will simulate a microfluidic water cycle. Pure water from the bulk solution will evaporate at the hot side and condense on the cold surface. These water droplets will grow in size and fall back into solution. The evaporation, water condensation and re-entry into the bulk solution leads to decrease (evaporation) and increase (rain fall) of the interface height. We therefore sought to determine how such wet-dry cycles and water precipitations would affect the coacervate droplets.

To do this, a dispersion of coacervate microdroplets (CM-Dex:PDDA molar ratio 6:1, [carboxyl]/[amine] = 5, total polymer concentration 20 mM, 10 mM Tris pH 8, 4 mM MgCl_2_, doped with 0.1 % FITC-labeled CM-Dex) was loaded into the thermal trap with a temperature gradient (hot side 34 °C, cold side 15 °C). Time-resolved optical microscopy images (Figure 4b and supplementary movie 3) showed that coacervate droplets accumulated at the gas-water interface and stacked to the warm surface of the trap as the height of the interface decreased from water evaporation. This had the effect of driving the accumulated coacervates into a quasi-dry state on the surface. The dry polymers (see arrow in Figure 4b) were later re-hydrated and the perturbative fluxes induced by the water precipitation led to their fragmentation. The resulting smaller daughter droplets fell into the bulk and circulated with the convection flow. These results show that water cycles can drive the fragmentation and fission of coacervate droplets. Again, additional experiments with CM-Dex:pLys mixtures showed that this process is general and can also take place when different types of coacervates are used (supplementary section 6).

Despite this, fission events were rarely observed. Out of a total of fifty-three experiments (average duration of ~2 hours each) which explored different polymer types, polymer concentrations, temperature gradients, buffers and trap geometries, we observed twelve division events. Of these twelve events, ten of them consisted of division by fragmentation (the type of Figure 4b). Two of them were of the type shown in Figure 4a. However, the division events may be happening more since we only image one of the many gas bubbles that were present in the chamber. Thus our count of droplet fissions may be underestimated. It is also important to note that our imaging protocol projected the view of the thermal trap on a 2D plane, and was therefore not able to distinguish objects or observe any dynamics in the perpendicular axis. In supplementary section 7, we thoroughly analyzed the experiment shown in Figure 4a to rule out possible artifacts deriving from the imaging.

Taken together, our results show two mechanisms by which the out-of-equilibrium behaviour induced by the thermal gradient at the gas-water interface of a microfluidic pore can drive droplet fission. This represents a viable route to coacervate fission and subsequent evolution within the prebiotically plausible scenario of a thermal pore.

Furthermore, to determine how robust the behaviour within the pore was, we characterized the effect of different temperature gradients (ΔTs between 10 and 60 °C), trap thicknesses (between 127 and 500 μm), and the volume of the gas bubbles (between 0.005 to 50 mm^3^) on dispersions of coacervate droplets. Within these broad range of conditions, the features of coacervate accumulation, fusion, wet-dry cycles and divisions were observed. It appears that differences in these three parameters can affect the sedimentation and accumulation properties, fusion and division events and the quantity of dried polymers on the surface of the pore. For example, steep temperature gradients induce a fast convection in the bulk which prevents sedimentation and induces a fast capillary flow that promotes the fusion between the droplets. The increased wet-dry cycles also promote the division mechanism by fragmentation (Figure 4b). On the other hand, droplet division by stretching would benefit from shallower temperature gradients, because the droplet needs to be slowly stretched in order to divide (Figure 4a). In addition, steep temperature gradients will affect the size and frequency of water precipitations and, consequently, the extent at which the gas-water interface moves up and down during the evaporation / water condensation cycles that affect the quantity of dried polymers.

In summary, the general properties of accumulation, fusion and division, drying and coacervate reentry are observed across a broad range of experimental conditions such as temperature gradient, the chamber thickness and the gas to liquid ratio. Tuning these experimental parameters will tune the dynamic behavior of the droplets in the pore. This provides exciting and plausible evidence that our observed phenomena of flow induced droplet maintenance, accumulation, fusion and fission could have taken place within rocky environments of early Earth, which had pores of different sizes, incorporated bubbles of different dimensions and were subject to different thermal gradients.

### Separation and selection of coacervate phenotypes

So far, we have determined the effect of the thermal trap with gas bubble on coacervates prepared from modified sugars, peptides and synthetic polymers. Despite the fact that PDDA was unlikely as a prebiotic molecule, we observed the general phenomena of accumulation, fusion, maintenance and fission by different mechanisms which appear independent of the chemical properties of the coacervate (Figure S2).

Recent studies have shown that compartmentalization by coacervation^12,37^ or the hydrophobic effect with fatty acids^38^ could complement the RNA world hypothesis by providing means to accumulate RNA and regulate RNA activity. Therefore, we wanted to determine the effect of the out-of-equilibrium dynamics of the thermal trap on dispersions of CM-Dex, pLys and RNA. To do this, dispersions of CM-Dex and pLys (molar ratio 4:1, [carboxyl]/[amine] = 7) with and without RNA (51 nt single-stranded, Figure 2e) were prepared at concentrations of 1.5 mM, 0.5 mM and from 0-5 μM respectively in 10 mM Tris pH 8, 4 mM MgCl_2_. In order to study the co-localization between RNA, CM-Dex and pLys, dual-channel fluorescence imaging was used. RNA was labelled with ROX (Carboxy-X-rhodamine) while 0.1 % of the coacervate components (CM-Dex or pLys) contained a FITC label (see Figure 5a). The microscope was equipped with an image splitter (Optosplit ii) containing the filterset for FITC and ROX to enable dual-channel fluorescence imaging.

After loading the dispersions of CM-Dex, pLys with RNA into the sample chamber, dual-channel fluorescence imaging showed that pre-formed small coacervate droplets (size < 15 μm) in the bulk colocalized RNA. Microscopy images showed that already prior to the thermal gradient, the droplets were rich in RNA and pLys with a weak signal attributed to CM-Dex. This indicates that RNA strongly competes with CM-Dex to form droplets with pLys. Indeed, thermophoretic measurements to obtain the binding constants between RNA with pLys and CM-Dex with pLys confirmed a higher affinity of RNA to pLys compared to CM-Dex (see supplementary section 8). Fitting to the dose response curve, we found that the K_D_ of the RNA:pLys complex (K_D_ < 11 nM) is an order of magnitude lower than the K_D_ of the CM-Dex:pLys complex (120 nM < K_D_ < 400 nM). This difference in K_D_ may be attributed to the fact that RNA has a higher charge density compared to CM-Dex. Therefore, whilst there is a small amount of CM-Dex within the droplet, CM-Dex will also be free in the coacervate dispersion. Upon inducing a thermal gradient (hot side 34 °C, cold side 15 °C), we observed the same phenomena as described previously i.e. that coacervate droplets accumulate at the interface and fuse together. Interestingly with the three coacervate components, dual fluorescence imaging of dispersions containing either FITC-labelled CM-Dex or pLys, with ROX labelled RNA showed that the droplets at the interface were larger and contained all three components (CM:Dex, RNA and pLys) (Figure 5a-c and supplementary movie 4) whilst the droplets in the bulk remained small and rich in RNA and pLys (Figure 5d-f). This observation is most likely due to the ability of the thermal trap to drive a strong accumulation of the RNA, pLys and CM-Dex in solution to the gas-water interface and induce an enrichment of the three components within the coacervate droplets, overcoming the equilibrium binding constants (supplementary section 8). Merging of the optical images shows that the microdroplets in the bulk have an overlap of the fluorescence signals of RNA and pLys (Figure 5d-f, supplementary section 9).

These results are important as they show that the thermal pore can generate and select for two different populations of coacervate droplets with different chemical compositions at the gas-water interface and within the bulk solution, which has not been previously reported upon.

We quantified the droplet size at the interface after applying the thermal gradient for 1 hour using the methodologies already described and as a function of RNA concentration. We observed that the final size of the coacervate protocells at the gas-water interface is inversely affected by RNA. In the presence of RNA, the average coacervate size dropped from 69 ± 31 μm down to 25 ± 9 μm (Figure 5g-j and Figure S8). As already shown in other studies^39^, a higher charge density can lead to the formation of smaller coacervate droplets. This is in fact what we observed, and we believe that the effect is driven by the stronger binding of RNA with pLys compared to CM-Dex with pLys.

The results show how the thermal trap can keep the coacervate droplets in a non-equilibrium state enabling energetically unfavorable interactions at the interface. This permits the formation and selection of two different populations of droplets within the pore with different physical properties and different compositions. We also show that the chemical composition of the coacervate droplets will affect their phenotype with smaller droplet size for increasing RNA concentration.

**Figure 5.**
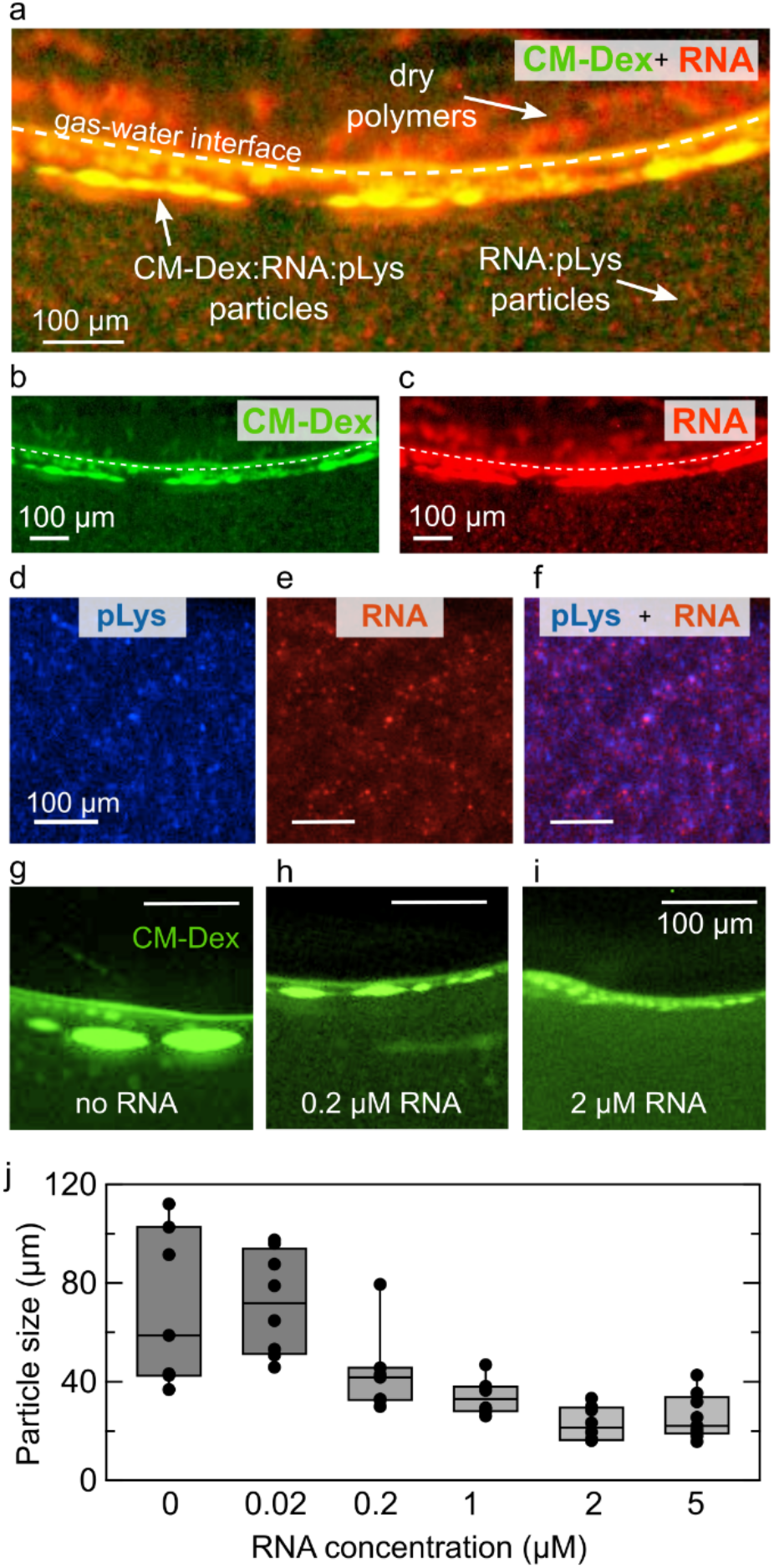
The thermal trap creates and separates two populations of coacervate droplets. a) Dual-channel fluorescence images of the CM-Dex:pLys:RNA coacervates in the thermal trap. CM-Dex and RNA were differentially labeled with FITC 0.1% and ROX 100%, respectively. The single pictures of the composite (a), are shown in (b) and (c), respectively. Small droplets (< 15 μm) enriched in RNA and pLys are formed in the bulk. Droplets enriched of all three components form instead at the gas-water interface. d) pLys channel (0.1% FITC-labelled), RNA channel (e) and composite image (f), showing co-localization between RNA and pLys in the bulk droplets. (g) no RNA, (h) RNA 0.2 μM and g) RNA 2 μM showing the droplets at the gas-water interface (CM-Dex fluorescence). h) Quantification of the size of CM-Dex:pLys droplets as a function of RNA conc. The bars indicate the average size and standard deviation of approximately 9 different coacervate droplets.

## Discussion

We showed that experimental primordial conditions - a millimeter-sized pore in a temperature gradient with a gas bubble - imparted specific selection pressures on dispersions of coacervate microdroplets. The thermal gradient across the pore drove a convection flow within the bulk solution and instigated the accumulation and growth of the coacervate droplets by fusion at the gas-water interface. The forces in the heated rock-like pores hindered the sedimentation of the coacervate droplets and the formation of large coacervate macrophases whilst permitting the maintenance of cell-like sized coacervate microdroplets for longer times. These droplets were elongated due to convection and capillary forces and underwent division after deformation at the gas-water interface. In addition, we observed division as a consequence of a water cycle within the gas bubble. The water precipitations induced the division and fragmentation of the coacervate material accumulated on the surface of the pore. These features were not observed in thermal traps in the absence of gas bubbles or at isothermal temperature indicating that this was a unique property of the thermal gradient and the gas bubble. This is the first example of the accumulation, fusion, maintenance and fission of coacervate protocells. We showed that this is a general phenomenon as we observed the same processes in coacervates with different chemical compositions and buffer conditions. These results represent a possible mechanism for the growth and division of membrane-free protocells on primordial Earth.

We also showed that the K_D_ determined the affinity of polyelectrolytes to form coacervates where oligonucleotides (RNA) had a higher propensity to form coacervates with polypeptides (pLys) compared to modified sugars (CM-Dex). The coacervate microdroplets that we studied seemed to be selective towards RNA incorporation, a molecule which can be catalytic. In an origin of life scenario, this process could give a selective advantage in terms of catalysis within a pool of coacervate protocells. The thermal trap generated two different populations of coacervate droplets, where droplets poor in CM-Dex were maintained in the bulk solution whilst CM-Dex rich droplets formed and accumulated at the gas-water interface. This finding shows that the environment of a thermal trap with a gas bubble enables energetically unfavorable coacervate droplets to form by driving the system into an out-of-equilibrium state. As a consequence, the thermal trap was able to generate and contain populations of coacervate droplets which differ in the chemical composition and size and therefore physical properties. In the presence of active RNA these genotypic and phenotypic differences would most likely lead to different activities within the droplet. The droplets at the gas-water interface would benefit from additional variability and non-equilibrium properties: preferential enrichment of longer oligonucleotides^28^; enhanced strand separation at lower temperatures^27^ (and therefore, lower hydrolysis rates); enhanced RNA catalysis induced by the presence of an additional polyanionic component that could lead to the change in material properties and the diffusion and reactions rates of RNA within the coacervate^40^.

This has important implications for demonstrating how thermal fluxes could have driven an evolutionary selection pressure on coacervate microdroplets giving experimental evidence for a key role within the origin of life scenario. In conclusion, our work shows that a temperature gradient with a gas bubble generates a unique environment for the accumulation, fusion, fission and selection of coacervate microdroplets. To the best of our knowledge this is the first time that these characteristics have been made accessible by physical forces alone - without chemical complexity or sophisticated machineries as seen in modern biology. This makes the gas bubble within a heated rock pore a compelling scenario to drive the evolution of membrane-free coacervate microdroplets on early earth.

## Materials and methods

CM-Dex sodium salt (10-20 kDa or 150-300 kDa, monomer: 191.3 g/mol), pLys hydrobromide (4-15 kDa or 150 kDa, monomer: 208.1 g/mol), and PDDA chloride (8.5 kDa, monomer: 161.5 g/mol), FITC-labelled pLys (15-30 kDa), FITC-labeled CM-Dex (15 kDa or 150 kDa), ATP (507.2 g/mol) were purchased from Sigma Aldrich Germany and used without further purification. Stock solutions of each of the coacervate components were prepared to a concentration of 1 M in milliQ water and stored at −20 °C until further use. RNA oligonucleotides were purchased from biomers.net Gmbh, with HPLC purification and re-dissolved to a final concentration of 100 μM in nuclease-free water. The sequence was (51 bases): 5’-UUA GCA GAG CGA GGU AUG T^ROX^AG GCG GGA CGC UCA GUG GAA CGA AAA CUC ACG. Every RNA strand was labeled with a ROX molecule (Carboxy-X-Rhodamine) positioned centrally in the sequence attached to the backbone of a Thymine and stored in pure nuclease-free water at a concentration of 100 μM.

The experiments were undertaken in a thin layer of PTFE (250 μm), which was cut with a defined geometry and then placed between a transparent sapphire and a copper back plate. The geometry of the PTFE sheet was designed to induce the incorporation of gas bubbles as shown in previous work^27,28^. The sapphire was in contact with a copper placeholder which was heated with rod resistors. The copper back plate was attached to an aluminum holder which was cooled with liquid water from a water bath (300F from JULABO). Temperature sensors (GNTP-SG from Thermofühler GmbH) were attached to the copper back plate and to the copper sapphire-holder to measure the outer temperatures of the cold and warm sides. The inner temperatures of the chamber were then calculated numerically based on the outer temperatures, the heat conductivities of the materials (coppers, silicon and sapphire) and their thickness. The outer warm target temperature was maintained constant via a PID loop implemented in LabVIEW, in order to control the output voltage to the rod resistors. The accuracy of the target temperatures was of ± 1 °C. The temperature differences that we used in the experiments shown here range from 15 to 30 °C.

Coacervate components were mixed together to the final desired concentration (2 to 20 mM) and immediately loaded into the microfluidic chamber. Dispersions of coacervates were prepared from either CM-Dex:PDDA or CM-Dex:pLys or CM-Dex:pLys: RNA in either 0.1 M Na+ Bicine buffer (pH 8.5) or 10 mM Tris and 4 mM MgCl_2_ (pH 8.0). The chamber was then loaded onto a fluorescence microscope (see supplementary section 1), focused on the cold wall and images were taken every 1 to 10 seconds for arbitrary time (usually 1 to 2 hours) using custom-built software LabVIEW.

Imaging was performed with a standard custom-built fluorescence microscope, equipped with a blue LED (470/29 nm), an amber LED (590/14 nm), excitation filters (482/35 nm, 588/20 nm), a dualband-pass dichroic mirror (transmission edges at 505 nm and 606 nm), a 5X objective and an image splitter containing a longpass filter (600 nm) and emission filters (536/40 nm, 630/50 nm). This filterset allowed for the imaging of FITC (Fluorescein isothiocyanate) and ROX (Carboxy-X-Rhodamine) respectively. The crosstalk between the channels was calculated following a standard protocol^27^ (supplementary section 1). A Stingray-F145B ASG camera (ALLIED Vision Technologies Gmbh) was used to acquire images. The voltage to the LEDs and the camera were controlled with the software LabVIEW (a scheme of the microscope is shown in figure S1). Image analysis of the droplets were undertaken using ImageJ or LabVIEW. The raw data from the two different illumination channels were merged together to generate the composite dual fluorescence image.

## Acknowledgements

Financial support came from: the European Research Council (ERC Evotrap, Grant Number 787356), the Simons Foundation (Grant Number 327125), the Quantitative Biosciences Munich Graduate School (QBM), MaxSynBio Consortium (jointly funded by the Federal Ministry of Education and Research (Germany) and the Max Planck Society), the MPI-CBG, the Cluster of Excellence Physics of Life of TU Dresden, EXC-1056 and the VW foundation (grant number 94743). We thank Lorenz Keil for sharing his expertise in the preparation of the setup for imaging and for programming support.

