## Supplementary information for "Non-equilibrium conditions inside rock pores drive fission, maintenance and selection of coacervate protocells"

**Supplementary information table of content:**

1. **Microscope scheme**
2. **Effect of buffer and polymer composition on coacervate assembly at the gas-water interface**
3. **Relationship between droplet growth and total polymer concentration**
4. **Control experiment without gas-water interface**
5. **Sedimentation of coacervate polymers**
6. **Division and fragmentation of CM-Dex:pLys coacervate droplets**
7. **Figure 4a extended: division by droplet stretching**
8. **Measurement of the binding constant of pLys:CM-Dex and pLys:RNA**
9. **Component ratio of the coacervates in the thermal pore**
10. **Figure 5 extended: RNA concentration vs final coacervate size**
11. **List of attached files**
12. **Author contribution**
13. **References**

### 1. Microscope scheme

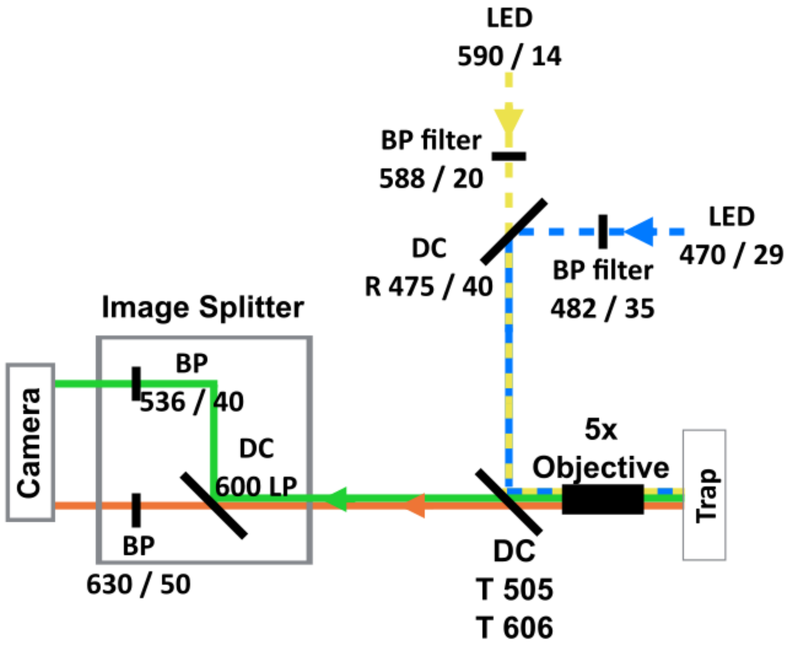


**Figure S1. Schematic of the custom-built microscope.** *The microscope was equipped with a blue LED (470/29 nm), an amber LED (590/14 nm), excitation filters (482/35 nm, 588/20 nm), a dual bandpass dichroic mirror (transmission edges at 505 nm and 606 nm), a 5X objective and an image splitter containing a longpass filter (600 nm) and emission filters (536/40 nm, 630/50 nm). This filterset allowed for the imaging of FITC (Fluorescein Isothiocyanate) and ROX (Carboxy-X-Rhodamine). A Stingray-F145B ASG camera (ALLIED Vision Technologies Gmbh) was used to acquire images. The numbers next to the filters and the LEDs correspond to the light wavelength in nanometers / FWHM.*

The crosstalk from the FITC channel to the ROX channel was calculated with the following standard protocol[1]:

$$crosstalk = \frac{DA_A}{AA_A}$$

where the first capital letter indicates the excitation wavelength (D), the second capital letter indicates the emission wavelength (A), and the subscript index indicates which dye was used (D = FITC, A = ROX). The crosstalk from the FITC to the ROX channel was measured to be approximately 7%. This is a relatively low value which would not have affected the qualitative comparison made in Figure 5.

### 2. Effect of buffer and polymer composition on coacervate assembly at the gas-water interface

We studied the effect of the buffer composition and the type of coacervate polymer on the assembly and growth of the coacervate droplets at the gas-water interface. Following the same procedure of the experiments shown in Figure 3, we ran several experiments with slightly different conditions of buffer or polymer composition, and temperature gradients. The mixtures were doped with 0.1 % FITC-labeled CM-Dex or ATP. The results are schematized in Figure S2. The droplet size over time was quantified and shown in figure 3e.

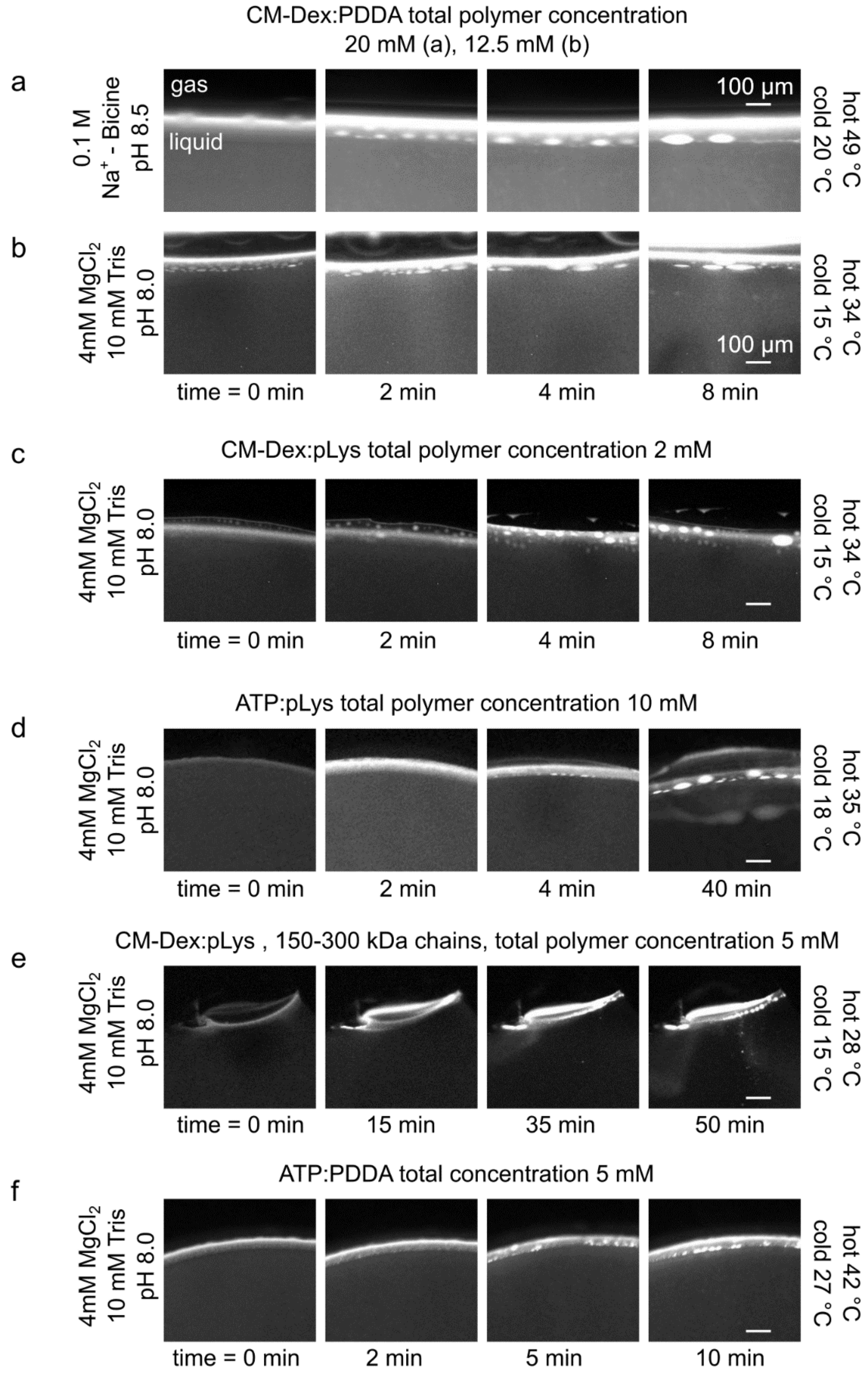


***Figure S2. Effect of the buffer and polymer composition on coacervate growth at the gas-water interface.*** *Growth mechanism by fusion for different coacervate types (indicated above the image), for different buffers (indicated at the left) and different temperature gradients (indicated at the right). The scale bar indicates 100 µm.*

When using a different buffer ($Na^+$-bicine or Tris-$MgCl_2$) or a different polymer (CM-Dex, pLys, PDDA or ATP), we observed only minor differences in the coacervate assembly process. The growth process (by fusion with other coacervate droplets), occurred with similar timescales and reached a similar threshold size in all the cases that we explored.

### 3. Relationship between droplet growth and total polymer concentration

We studied the effects of the total polymer concentration on the growth of the coacervate. To this end, we performed a series of experiments in a thermal gradient (hot side 49 °C and cold side 20 °C), buffer (4 mM $MgCl_2$, 10 mM Tris, pH 8.0) and the CM-Dex: PDDA (molar ratio 6:1, [carboxyl]/[amine] = 5), varying the total polymer concentration between 1 and 20 mM, doped with 0.1 % FITC-labeled CM-Dex. After inserting the coacervate solution in the thermal trap, we took a microscopy image every ~ 1 second. The images were analyzed with ImageJ or LabVIEW to determine the growth rate and the average droplet size at the interface. The results are shown in figure S3.1. Every data point in the plot corresponds to an independent experiment.

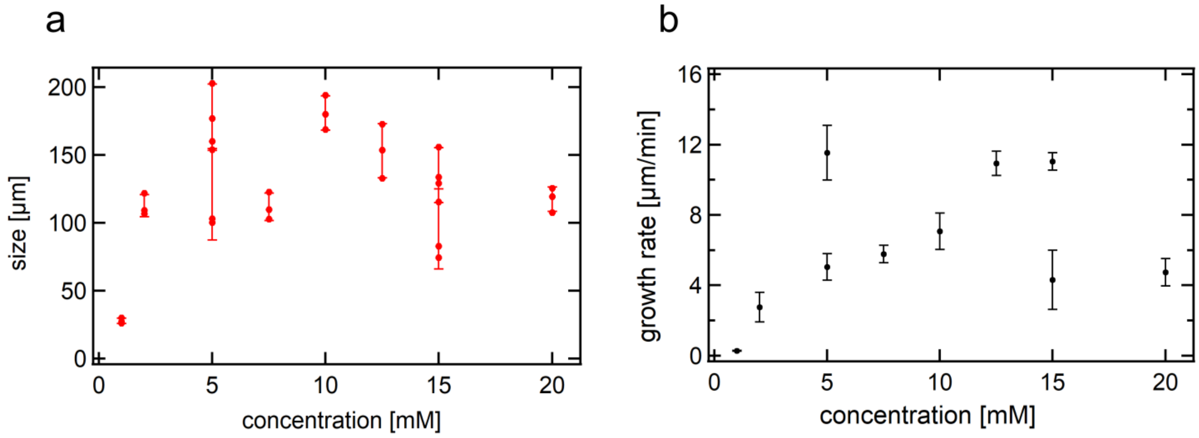


***Figure S3.1. No clear relationship between coacervate growth and total polymer concentration.*** *(a) Average size at steady state (after ~ 1h of thermal gradient). Every set of data is represented by 3 points (the 3 largest coacervate droplets observed in the experiment) and the error bar (standard deviation). b) Growth rate of the coacervate droplets as a function of polymer concentration. Error bars correspond to the standard deviation of the growth rate extracted from the linear fit of the size vs time traces (here not shown).*

Due to the intrinsic noise of our experiments, we could not determine any significant effect of the total polymer concentration on the final droplet size at steady state and their growth rate.

Below the critical coacervate concentration (CCC), polymers are not able to coacervate and form droplets. We tested whether the accumulation properties of the heated gas-water interfaces in our pores were able to concentrate the polymers to cross the CCC threshold and trigger coacervation. We used a CMDex:PDDA mixture (molar ratio 6:1, [carboxyl]/[amine] = 5), with a total polymer concentration below the CCC, doped with 0.1 % FITC-labeled CM-Dex. In previous experiments the CCC of CM-Dex:PDDA >1mM. Therefore, in the experiments that follow, we used a total polymer concentration of 0.2 mM and 0.05 mM, in a buffer containing 4 mM $MgCl_2$ and 10 mM Tris at pH 8 with a temperature gradient (hot side 49 °C, cold side 20 °C) (below the CCC). The polymer accumulation at the interface was imaged with fluorescence microscopy (Figure S3.2.).

Our results show that no coacervate droplets were observed in the system but there was visible polymer accumulation at the gas-water interface. This observation suggests that thermal trap acts at the mechanical level to favor the assembly and fusion of the existing coacervate droplets.

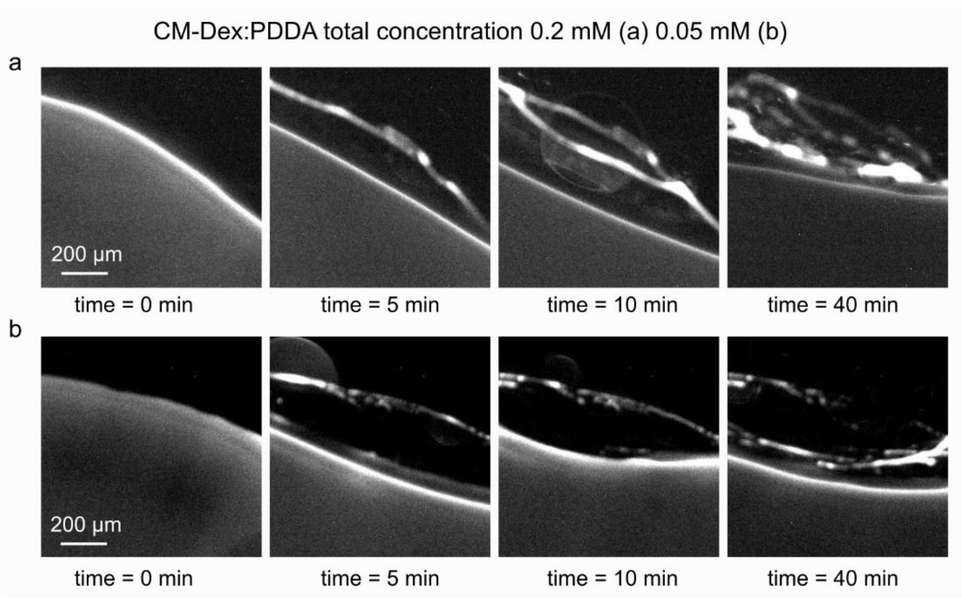


***Figure S3.2. No coacervation below the critical coacervate concentration.*** *Optical microscopy images of the gas-water interface in the thermal trap at different times (0, 5, 10 and 40 minutes) of the coacervate mixture in the thermal pore, for 0.2 mM (a) or 0.05 mM (b) total polymer concentration. No coacervate droplets were observed at the gas-water interface or in the bulk despite the visible polymer accumulation.*

### 4. Control experiment without gas-water interface

In order to see the effect of the thermal gradient on coacervation, we characterized the effect of a thermal flow on coacervate droplets in a control experiment without gas bubbles (Figure S4).

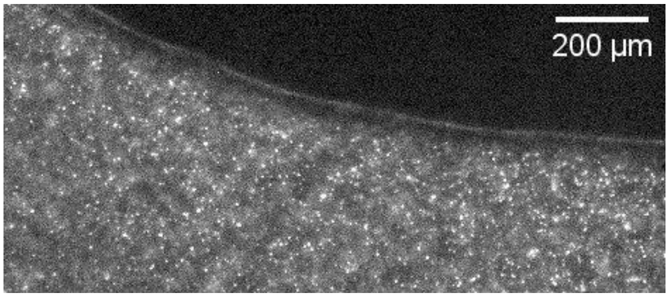


***Figure S4. No large coacervate droplets in a thermal gradient without a gas bubble.*** *Coacervate droplets were prepared from CM-Dex:pLys (molar ratio 4:1, [carboxyl]/[amine] = 7, total polymer concentration 2 mM), buffer 10 mM Tris, 4mM $MgCl_2$, pH 8.0, temperature gradient of 29 °C. The image shows the fluorescence of pLys (0.1 % labeled with FITC). Small coacervate droplets (size < 10 µm) are seen moving with the convection flow. No larger coacervates (size > 15 µm) were observed forming in the system.*

In the absence of the gas bubble, no large coacervates form in the chamber. Small coacervate droplets (< 15 µm), which spontaneously formed during the initial mixing of the polymers, could be seen in the bulk transported by the convection flow at a speed of about 1.6 ± 0.4 µm/s. However, they did not fuse to form larger droplets. The gas-water interface has the role of accumulating small microdroplets and promote their fusion. In the experiments with a gas-water interface, we observed that the coacervate droplets could grow up to an average size of 150 µm. Sometimes we could observe coacervate droplets as large as 300 µm.

### 5. Sedimentation of coacervate polymers

We characterized the behaviour of the coacervate droplets under equilibrium condition. In the absence of a thermal gradient, no convection occurs in the liquid phase, and the coacervate droplets fall to the bottom of the channel, driven by gravity, to form a single coacervate macrophase. We studied the sedimentation dynamics of CM-Dex:pLys (molar ratio 4:1, [carboxyl]/[amine]=7, total polymer concentration 8 mM in 4 mM $MgCl_2$, 10 mM Tris at pH 8.0, doped with 0.1 % FITC-labeled CM-Dex) in a microfluidic chamber at room temperature (22 °C, no temperature gradient). The droplet size was determined as its horizontal width. The results are shown in figure S5.1. The growth rate of the coacervate droplets by sedimentation is much slower (requires hours) than the growth rate at the heated gas-water interface shown in figure 3e (requires minutes). On the other hand, the final droplet size is much greater, and only constrained by the size of the microfluidic chamber.

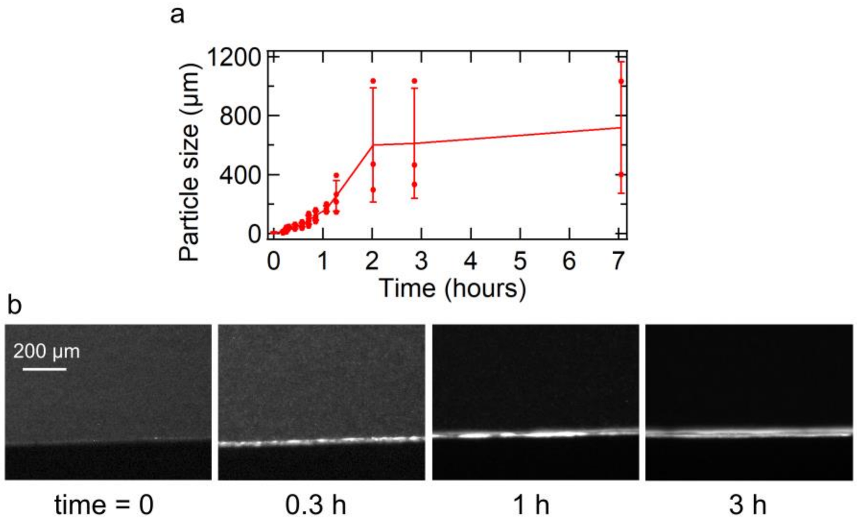


***Figure S5.1. Sedimentation of coacervate droplets induced by gravity.*** *a) Quantification of droplete size over time. The droplets slowly sediment at the bottom of the chamber and fuse to become larger over the course of several hours. Error bars indicate standard deviation (n~3). b) Snapshots of the sedimentation at different times. The fluorescent layer at the bottom is the site of sedimentation.*

In the presence of the thermal gradient, a circular convective flow in the bulk arises. As we have already seen in our experiments, it transports the small coacervate droplets around in the bulk, preventing the sedimentation.

We simulated the sedimentation process by a 2D finite element simulation, using the software Comsol Multiphysics. We modeled the heat transfer, laminar flow, diffusion and transport of coacervate droplets in a thermal gradient, in a chamber equivalent to the one we used in our experiments (5 mm x 500 µm). The density of the coacervate droplets was set between 1.01 and 1.10 g/cm$^3$ and their radius was set to 5 µm as determined from our experiments. The total polymer concentration was set to 1 mM. The diffusion coefficient was calculated from the radius and the density, using the following Stokes-Einstein equation:

$$D = \frac{k_B\, T}{6\, \pi\, r_0\, \mu_B(T)}$$

where $k_B$ is the Boltzmann constant 1.38e-23 J/K, $T$ is the temperature, $r_0$ is the droplet radius ad $\mu_B$ is the bulk viscosity. The thermal gradient was varied between 0 °C and 85 °C. Results are shown in figure S5.2.

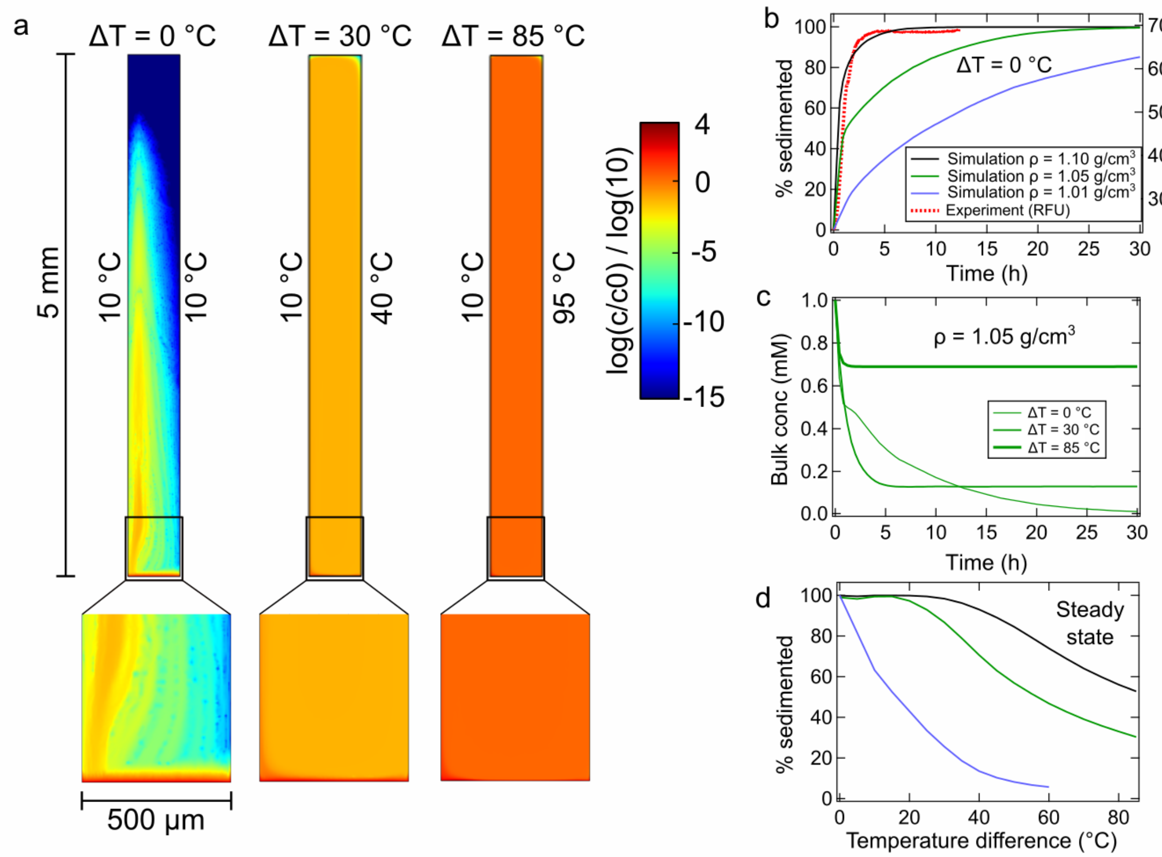


***Figure S5.2. Finite element simulation of the sedimentation of coacervate droplets in our thermal trap.*** *a) Distribution of coacervate concentration in a trap section after 10 h, for ΔT of 0, 30 and 85 °C. The coacervate density coefficient used here was 1.1 g/cm$^3$. b) Sedimentation % over time at isothermal temperature for 3 different densities. Red dotted line corresponds to experimental data (the experiment illustrated in Figure S5.1). c) Bulk coacervate concentration over time calculated for different temperature gradients. d) Sedimentation % at steady state (simulation time = 30 h) as a function of the temperature gradient ΔT and for different densities.*

In the absence of the thermal gradient, almost 100% of the coacervate droplets sediment to the bottom of the chamber within 10 hours, driven by gravity (Figure S5.2a, left). The timescale of sedimentation depends on the density coefficient of the droplets. The best agreement between experiments and simulation occurs for a density coefficient between 1.10 and 1.05 g/cm$^3$ (Figure S5.2b). Note that the comparison between experiment and simulations is purely qualitative (expressed in arbitrary RFU), due to the difficulty in accurately calculating the total fraction of sedimented coacervates by fluorescence microscopy.

The total sedimented fraction is determined by the thermal gradient (Figure S5.2a middle, right pictures, and S5.2c-d). Our results show that a fraction of droplets will sediment even in the presence of the thermal gradient. However, our simulations show that temperature gradients between 19 and 40 °C will prevent the sedimentation of the coacervate fraction by 3 - 30 %. Furthermore, simulations, up to 96 hrs show that the droplets remain in the bulk solution indicating that with the thermal flow the droplets are prevented from sedimenting (data not shown). Note that the data shown in Figure S5.2d represent the final sedimented fraction at steady state.

Experiments undertaken within the thermal trap with dispersions of coacervate droplets (ATP:PDDA droplets, 5 mM total concentration) that were subjected to shallow temperature gradients (hot side 42 °C, cold side 27 °C) showed droplets at the gas-water interface after 30 hrs (Figure S5.3). Taking this into account, it is likely, that our experiment will reduce sedimentation further as the gas-water interface introduce additional flows to the system (e.g. capillary flows) that accumulate and maintain coacervate droplets at the gas-water interface. In summary, the thermal gradient is able to reduce the sedimentation of coacervates, maintaining a fraction of droplets to circulate in the bulk.

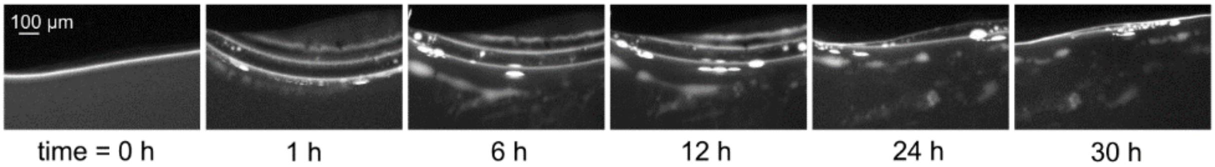


***Figure S5.3. Coacervate droplets are retained at the gas-water interface and do not sediment.*** *ATP:PDDA coacervate droplets formed at the gas-water interface do not undergo sedimentation for at least 30 hours.*

### 6. Division and fragmentation of CM-Dex:pLys coacervate droplets

We have already shown in figure 4 that CM-Dex:PDDA droplets in a thermal gradient can divide and be fragmented by the forces acting at the gas-water interface (capillary flows and perturbative fluxes). In additional experiments, we observed the same division and fragmentation processes happening for solutions containing a different coacervate composition: CM-Dex:pLys (molar ratio 4:1, [carboxyl]/[amine]=7, total polymer concentration 5 mM, doped with 0.1 % FITC-labeled CM-Dex). The results are shown in the Figure S6.

The coacervate droplet at the gas-water interface (yellow arrow) was stretched by the capillary forces, until the coacervate droplet started to divide, and the two daughter droplets separated. The event occurred in a time window of approximately 200 seconds.

Figure S6b shows the fragmentation of coacervates. The precipitation of water re-increased the water level and moved the gas-liquid interface slightly upwards. Therefore, the accumulated coacervates that were stuck in a quasi-dry state on the hot sapphire were re-dissolved into the liquid. The perturbative fluxes induced by the water precipitation induced the fragmentation of the polymers, creating smaller droplets that fell into the bulk and started circulating with the convection flow.

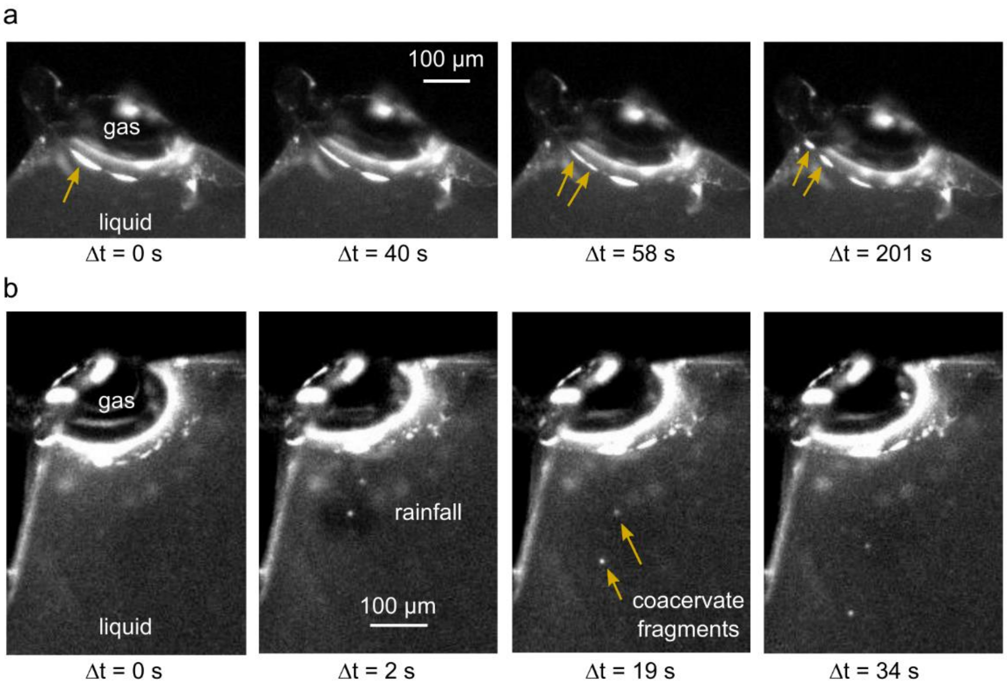


***Figure S6. Division of CM-Dex:pLys coacervate droplets at the gas-water interface.*** *a) Fission of a CM-Dex:pLys droplet into two smaller droplets induced by forces at the gas-liquid interface. b) Water precipitation rehydrates the stuck coacervates and induces fission by fragmentation. The temperature gradient for both experiments was: hot side 26 °C, cold side 15 °C.*

The division mechanisms described here and in figure 4 are purely physical phenomena. The chemical composition of the coacervate droplets seems to play a minor role, as we observed similar events for coacervate droplets made of CM-Dex:PDDA or CM-Dex:pLys. We therefore consider these division mechanisms to have played an important role in the primordial division of protocells, since they derive from the physical properties of the environment only and do not require any specific chemical component or active biological machinery. The simple and ubiquitous setting of a gas bubble within a thermal gradient contains all the physical properties to trigger these type of division mechanisms.

### 7. Figure 4a extended: division by droplet stretching

The dynamics of the dividing coacervate droplet (Figure 4a) was tracked throughout the timeframe of the experiment to confirm that the division event was not an artefact arising from the 2D imaging technique, as this type of imaging does not provide information in the x-axis of our thermal pore. In particular, we wanted to test whether two coacervate droplets could have hidden behind one other and then moved parallel in the x-axis, creating a visual artifact similar to a division event.

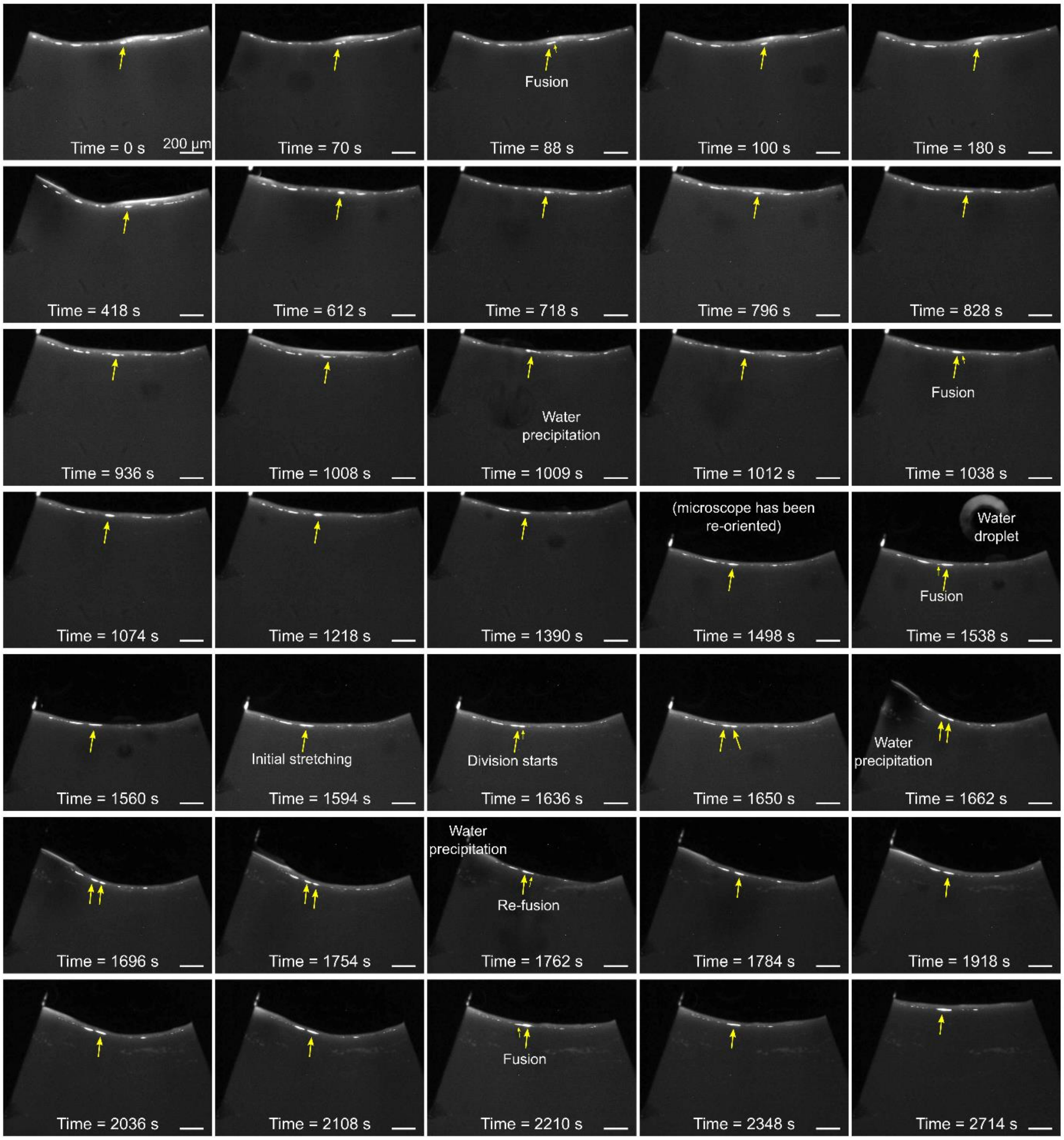


***Figure S7. Coacervate division by stretching.*** *Images at different times of the coacervate droplet that underwent division in Figure 4a. Experimental conditions were: CM-Dex:PDDA (molar ratio 6:1, [carboxyl]/[amine] = 5), total concentration 2 mM in 4 mM $MgCl_2$, 10 mM Tris, pH 8.0, doped with 0.1 % FITC-labeled CM-Dex. The temperature gradient was 19 °C with the hot side 34 °C, cold side 15 °C. The time reported in the pictures is not the absolute time of the start of the experiment, but is relative to the arbitrary starting point indicated in the first picture.*

Careful analysis of the full time frame (frame rate 0.75 fps) of coacervate division events at the air-water interface (Figure S7) showed repeated fusion and water precipitation events for many minutes, until the droplet started to undergo division (time ~ 1500s). We do not observe any indication of a second droplet moving behind the droplet of interest (indicated with a yellow arrow). It can be also seen that the droplet of interest remains distinct for several minutes before undergoing a division event. As the constrained thickness of the pore (x-axis size) is 250 µm and the daughter droplets measure approximately 90 µm, it is highly unlikely that two droplets of similar size will coexist behind each other without coalescence. Especially, when considering that capillary flow at the gas-water interface pushes towards the warm side that facilitates fusion events and that we typically observe fusion events between droplets in close proximity to each other within 10 seconds (Figure 3d). Our results indicate that there is no artefact of droplet fusion that arises from the imaging method used in our experiments.

### 8. Measurement of the binding constant of pLys:CM-Dex and pLys:RNA

In figure 5, we observed that RNA:pLys droplets are preferentially formed when a mixture of RNA, pLys and CM-Dex is provided. To gain a better understanding in the mechanism responsible of that phenomenon, we measured the binding constant $K_D$ of the RNA:pLys and of the CM-Dex:pLys complexes. Measurements have been done using a Nanotemper NT.115 Pico machine, which measures a binding dependent fluorescence signal upon local heating with IR laser.

Serial dilutions of CM-Dex (15 kDa) or RNA (single stranded, 51 nt) were mixed together with a constant amount of FITC-labeled pLys (15-30 kDa). While the final FITC-pLys concentration was maintained constant at 20 nM, CM-Dex and RNA concentrations spanned many orders of magnitude ranging from 0.1 nm to 1 µM. The solutions of CM-Dex: pLys or RNA: pLys were inserted into thin glass capillaries and placed on an alluminium holder and then inside of the Nanotemper NT.115 Pico machine. Therefore, the fluorescence of the sample was measured over time. A focused IR laser beam was used to locally heat the solution inside the capillaries, and the intensity response was measured. To calculate the $K_D$, the obtained data have been fitted with the following model:

$$\boldsymbol{f(Conc)} = \boldsymbol{U} + \frac{(\boldsymbol{B}-\boldsymbol{U})\cdot\left(\boldsymbol{Conc}+\boldsymbol{TargetConc}+\boldsymbol{K_D}-\sqrt{(\boldsymbol{Conc}+\boldsymbol{TargetConc}+\boldsymbol{K_D})^2-4\cdot\boldsymbol{Conc}\cdot\boldsymbol{TargetConc}}\right)}{2\cdot\boldsymbol{TargetConc}} \tag{1}$$

where *U* and *B* indicate the fluorescence response values of the unbound (after IR heating) and bound (no IR heating) states, respectively. TargetConc corresponds to the concentration of the labeled species (pLys), and $K_D$ corresponds to the binding constant. Conc indicates the concentration of CM-Dex (Figure S8.1a) or RNA (Figure S8.1b).

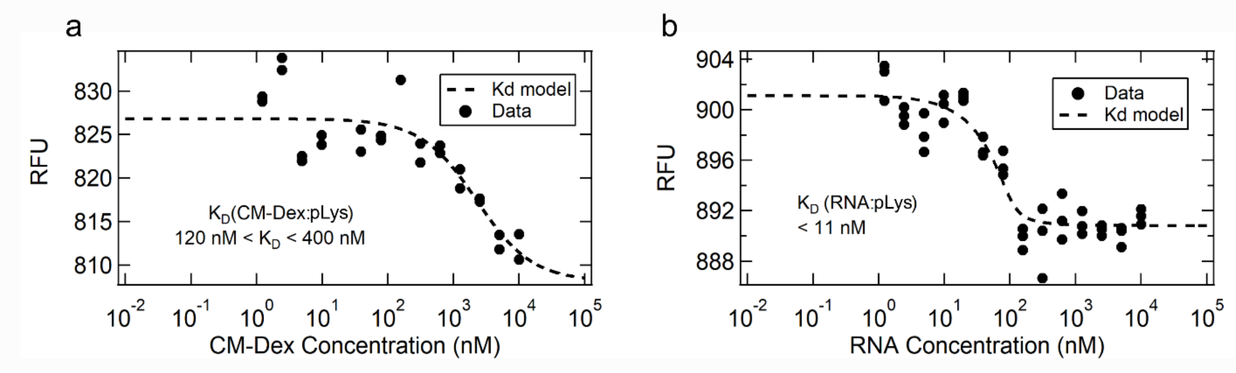


***Figure S8.1. Measurement of the binding constant $K_D$.*** *Dose-response curves for the CM-Dex: pLys complex (a) or for the RNA: pLys complex (b). The dots indicate experimental data, and the dashed line corresponds to the fit (equation 1).*

Instead of estimating a single value for the binding constant, we preferred to estimate a range where it most likely lies within. We achieved that by estimating the NRMSD (Normalized Root Mean Squared Deviation) between the data and the model. We arbitrarily chose a NRMSD threshold of 15% to estimate the range. NRMSD was defined as:

$$NRMSD = \frac{RMSD}{RFU_{max} - RFU_{min}}$$

where *RMSD* corresponds to the root mean squared deviation of the data. $RFU_{max}$ and $RFU_{min}$ are the maximum and minimum values of the RFU data. The binding constant ($K_D$) of the CM-Dex:pLys complex resulted to be higher than the $K_D$ of the RNA:pLys complex (120 nM <$K_D$ < 400 nM against $K_D$ < 11 nM), suggesting stronger binding of the RNA:pLys complex. This result can possibly explain what we observed in figure 5. In a solution containing all the three components (CM-Dex, RNA and pLys), pLys preferentially creates complexes with RNA, since their binding is stronger.

To further investigate whether the separation of coacervate populations in the thermal trap is driven by overcoming the binding constants at the gas-water interface, we performed additional control experiments. We studied a mixture of CM-Dex, pLys and RNA (total polymer concentration 2 mM) in the absence of a gas bubble, at temperature gradients (34 - 15 °C, as described previously) and at higher temperatures gradients (81 - 74 °C). The higher temperature is expected to overcome the binding constants, leading to the formation of bulk coacervate droplets enriched in all 3 components: the same effect induced by the gas-water interface (Figure 5).

At low temperatures (Figure S8.2a), the bulk coacervate droplets seemed to be mostly made of pLys and RNA with a low, almost indistinguishable fluorescence signal from CM-Dex within the coacervate droplets (Figure S8.2 left). In comparison, experiments undertaken at higher temperatures (Figure S8.2b) show evidence of three components (CM-Dex, pLys and RNA) by three fluorescent channels, suggesting that the bulk droplets were enriched in all three components. Our results indicate that, the higher temperatures could override the binding constant and trigger the interaction between all the three polymer components. Therefore, these results support our hypothesis that the gas-water interface helps to overcome the binding constants at lower temperatures.

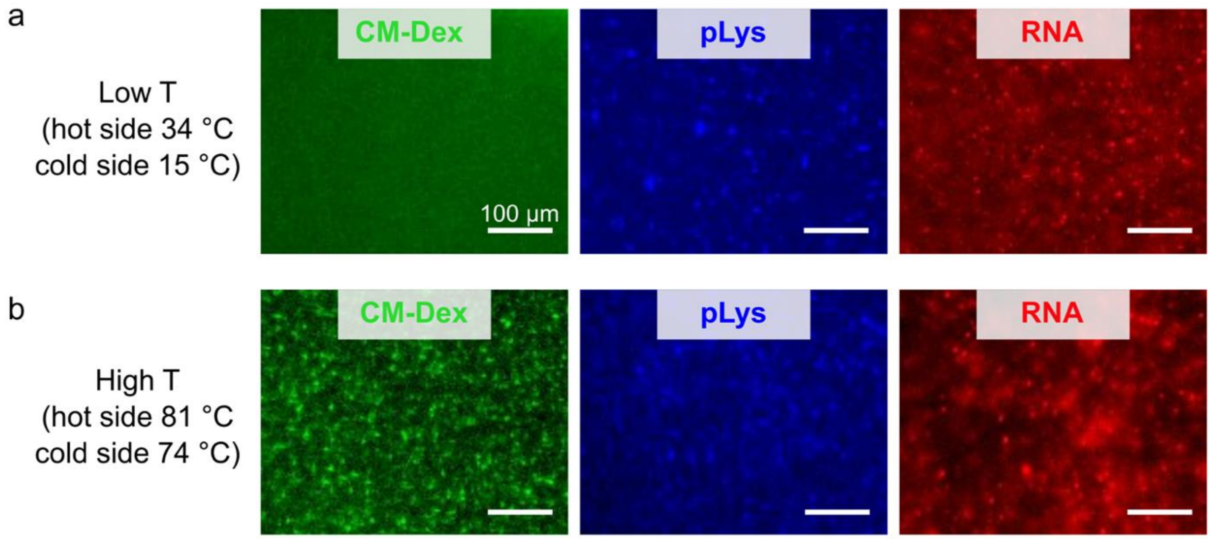


***Figure S8.2. Bulk droplets enriched in all three components form at higher temperatures.*** *a) At low temperatures (34 °C - 15 °C gradient), the bulk droplets are preferentially enriched in pLys and RNA, while poor in CM-Dex. b) At higher temperatures (81 °C - 74 °C gradient)*

*the binding constants are overridden by the increased kinetic energy and the bulk coacervate droplets are enriched of all three components. Every picture corresponds to a different experiment. The solutions contained: 10 mM $MgCl_2$, 10 mM Tris, pH 8. CM-Dex:pLys [carboxyl]/[amine] = 7, molar ratio 4:1. Total polymer concentration was 2 mM, RNA concentration 1 µM. Note that every picture corresponds to a separate experiment.*

### 9. Component ratio of the coacervates in the thermal pore

To characterize the effect of the water-air interface on properties of the coacervate droplets we measured the zeta potential (ζ) of coalescing coacervate droplets (CM-Dex:pLys) in bulk solution over time and compared this to any observed changes in the charge ratio determined by image analysis of the droplets within the thermal trap. The ζ potential corresponds to the difference between the potential on the shear surface of the droplet and the potential of the solution and is an indirect measure of the charge ratio at the surface of the droplet. To do this, dispersions of CM-Dex:pLys coacervate droplets at 2 mM in 10 mM Tris and 4 mM $MgCl_2$ at pH 8 (CM-Dex:pLys molar ratio 4:1, [carboxyl]/[amine] = 7) were prepared and immediately loaded into folded capillary cells (DTS1070) so that the water level was above the gold electrodes with no air bubbles observable to the eye. The cell was loaded into a Zetasizer Nano ZS (Pzen 5600) that was preheated at 30 °C. The sample was incubated for 10 minutes and 5 runs of 10 seconds were taken using the 173° backscatter mode to obtain dynamic light scattering data. This measurement was repeated 3 times. The sample was then subjected to zeta potential measurements which consisted of 10 secs equilibration time followed by 3 measurements of 10 runs with 300 secs between each measurement. This cycle was repeated over at least 2.5 hrs. The Zeta sizer was controlled using the manufacturers zetasizer software which undertook the analysis of the data using the general purpose analysis mode (light scattering) and the Smoluchowski model (Zeta measurements) using 1.378 and 1.334 for the refractive index for the coacervate and supernatant respectively.

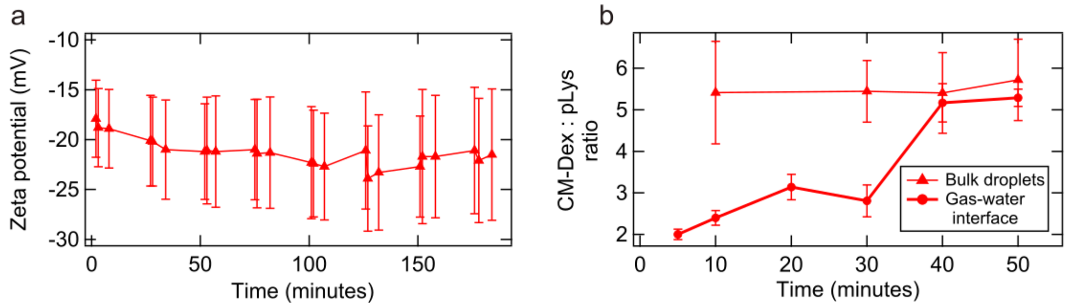


***Figure S9.1. Surface charge remains constant over time in the bulk droplets, while changes at the gas-water interface.*** *a) Measurement of the Zeta potential of the bulk droplets (isothermal bulk experiment). The error bars represent the standard deviation. b) Ratio between CM-Dex and pLys (no RNA) in the bulk droplets (triangles) or at the gas-water interface (circles). The experiment of (b) was made in a thermal trap (cold side 10°C, warm side 36 °C).*

We did not find a significant change in the surface charge over time (Figure S9.1a). Even though this measurement was performed in the bulk solution and under isothermal conditions, we observed analogous behavior within the bulk region in the trap experiment (Figure S9.1b). This was determined by fluorescence microscopy, by performing different experiments with FITC-labeled CM-Dex or pLys. We characterized the ratio of the polymers of the coacervate droplets assembled at the gas-water interface and in the bulk of our thermal pores. Conversely, at the gas-water interface, a change in the polymeric ratio could be observed during time. This could be attributed to differential accumulation of the individual polymers at the gas-water interface, driven by different diffusion coefficients, that variably enriched the droplets.

Furthermore, we undertook image analysis of dispersion of CM-Dex:pLys:RNA (51 nt) (1.6 mM, 0.4 mM, 1 µM, respectively), in a buffer containing 10 mM Tris, 4 mM $MgCl_2$, pH 8 within the thermophoretic pore. The temperature gradient was: hot side 34 °C, cold side 15 °C. The data shown in Figure S9.2 have been obtained by analyzing the dual-channel fluorescence movies over time where two of the three polymers were labeled. [Note that the ratios in Figure S9.2 are given as molar ratios between the polymers, not as ratios between chemical groups (e.g. [carboxy]/[amine]).

The analysis shows that the ratio of the components in the droplets changes over time. Both the ratios CM-Dex:RNA (Figure S9.2a) and pLys:RNA (S9.2b) decreased over time, while the CM-Dex:pLys ratio (S9.2c) remained almost constant. This can be explained by the fact that the droplets, both at the interface and in the bulk (Figure S9.3)., become enriched in RNA over time. The coacervate droplets in the bulk were particularly poor in CM-Dex in comparison to the coacervates assembled at the gas-water interface (Figure S9.2d).

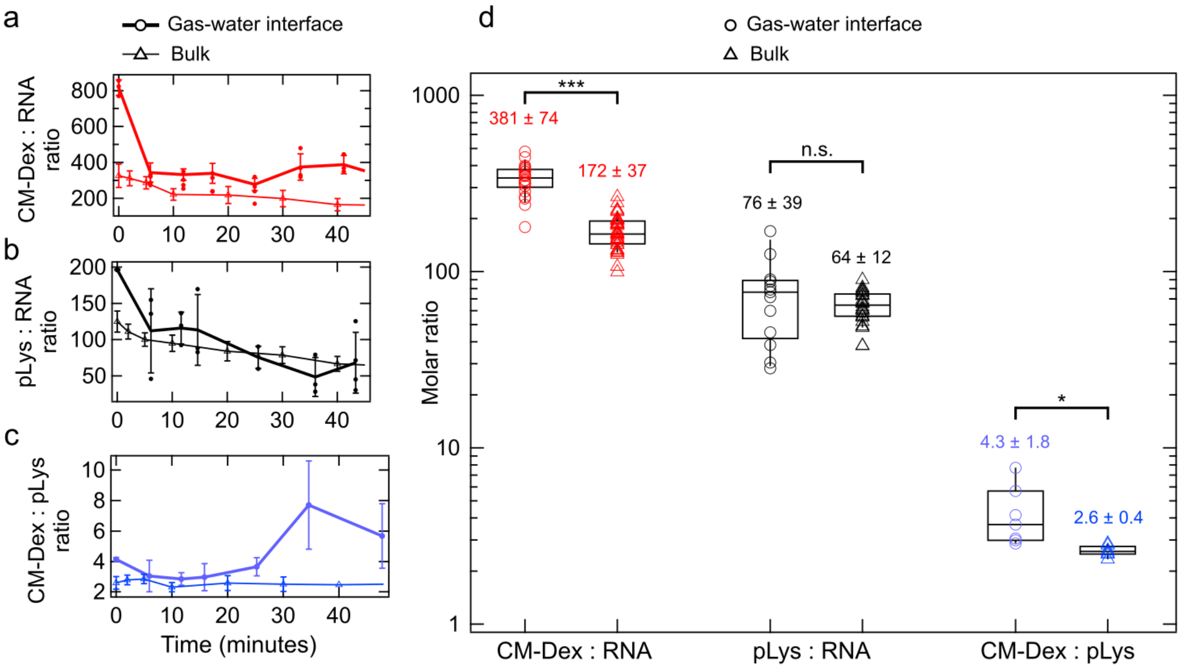


***Figure S9.2. Component ratio of a CM-Dex:pLys:RNA mixture in the thermal pore.*** *The droplets become enriched in RNA over time. CM-Dex:RNA (a), pLys:RNA (b) and CM-Dex:pLys molar ratios over time calculated on the coacervate droplets at the gas-water interface (thick lines with circles) or in the bulk (thin lines with triangles). d) Box plots of the ratios from a-c at steady state. The statistical difference between gas-water interface (circles) and bulk (triangles) was tested with a t-test.*

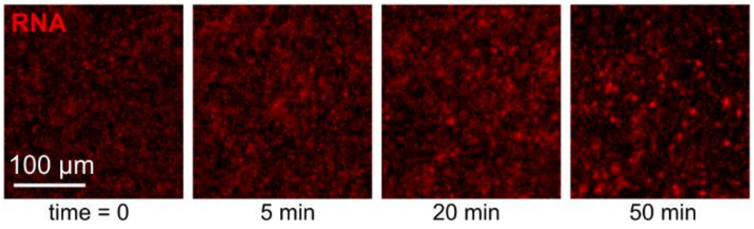

***Figure S9.3. RNA enrichment in the bulk coacervate droplets over time.*** *Fluorescence images (RNA fluorescence) of the bulk coacervate droplets at different times. An increase in the fluorescence levels is clearly visible.*

### 10. Figure 5 extended: RNA concentration vs final coacervate size

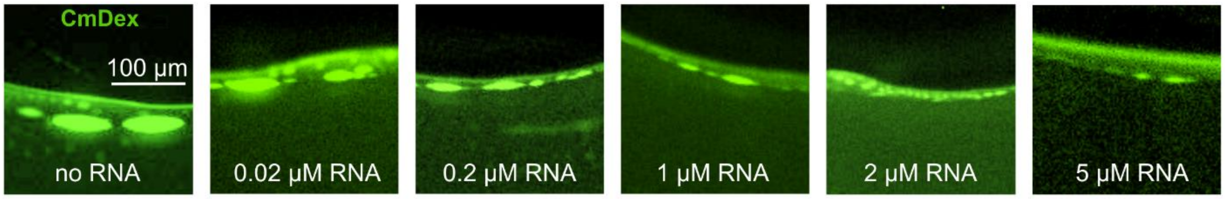


***Figure S10. RNA reduces the size of CM-Dex:pLys droplets in a concentration-dependent manner.*** *From left to right: no RNA, 0.02 mM RNA, 0.2 µM RNA, 1 µM RNA, 2 µM RNA, 5 µM RNA were added in a solution of CM-Dex:pLys (molar ration 4:1, [carboxyl]/[amine] = 7) at a concentration of 2 mM containing 0.1% of FITC-labeled CM-Dex in a buffer made of 4 mM $MgCl_2$, 10 mM Tris at pH 8. The images show the resulting coacervates at the gas-water interface after ~1h of temperature gradient (hot side 34 °C, cold side 15 °C).*

### 11. List of attached files

supplementary_movie_1_coacervate_fusion.avi
supplementary_movie_2_coacervate_division.avi
supplementary_movie_3_coacervate_fragmentation.avi
supplementary_movie_4_dual_channel.avi

### 12. Author contribution

A.I. and D.T. performed the experiments and analyzed the data. A.I., D.T., J.S., A.K., C. M., D.B. and T-Y D.T. conceived and designed the experiments. A.I, D.T., D.B and T-Y D.T wrote the manuscript.
